# Robust Inference of Bi-Directional Causal Relationships in Presence of Correlated Pleiotropy with GWAS Summary Data

**DOI:** 10.1101/2022.03.02.482630

**Authors:** Haoran Xue, Wei Pan

## Abstract

To infer a causal relationship between two traits, several correlation-based causal direction (CD) methods have been proposed with the use of SNPs as instrumental variables (IVs) based on GWAS summary data for the two traits; however, none of the existing CD methods can deal with SNPs with correlated pleiotropy. Alternatively, reciprocal Mendelian randomization (MR) can be applied, which however may perform poorly in the presence of (unknown) invalid IVs, especially for bi-directional causal relationships. In this paper, first, we propose a CD method that performs better than existing methods regardless of the presence of correlated pleiotropy. Second, along with a simple but yet effective IV screening rule, we propose applying a closely related and state-of-the-art MR method in reciprocal MR, showing its almost identical performance to that of the new CD method when their model assumptions hold; however, if the modeling assumptions are violated, the new CD method is expected to better control type I errors. Notably bi-directional causal relationships impose some unique challenges beyond those for uni-directional ones, and thus requiring special treatments. For example, we point out for the first time several scenarios where a bi-directional relationship, but not a uni-directional one, can unexpectedly cause the violation of some weak modeling assumptions commonly required by many robust MR methods. Finally we applied the proposed methods to 12 risk factors and 4 common diseases, confirming mostly well-known uni-directional causal relationships, while identifying some novel and plausible bi-directional ones such as between BMI and T2D, and between BMI and CAD.

## 1 Introduction

It is of great interest to infer causal relationships between pairs of complex traits or diseases such as for treatment/intervention development and drug repurposing [45, 10], which however is quite challenging and had barely been touched until recently. The availability of large-scale GWAS summary data and the use of SNPs as instrumental variables (IVs) in Mendelian randomization (MR) have made it possible for such inference [12, 13, 14]. Recently several methods based on comparing correlations between SNPs/IVs and each trait have been proposed for such a purpose, including Steiger’s method based on a single SNP (that is assumed to be a valid IV) [21], CD-Ratio and CD-Egger based on multiple SNPs, which can be more powerful than Steiger’s method [51]. CD-Egger, similar to Egger regression in MR [3], is also more robust than the other two methods by allowing invalid IVs under the InSIDE assumption; that is, CD-Egger allows invalid IVs with uncorrelated pleiotropy, but not correlated pleiotropy [32]. The first goal here is to develop a correlation-based causal direction (CD) inference method based on constrained maximum likelihood, called CD-cML, and show its higher power and robustness than the above methods, especially in the presence of correlated pleiotropy. Given the wide-spread pleiotropy [44, 47], it is of utmost importance for any method to be robust to pleiotropy, especially correlated pleiotropy that is more challenging to deal with.

As an alternative, reciprocal (also called bidirectional) MR can be applied by treating each of the two traits as the exposure while the other as the outcome [42, 50]. However, as shown in [34, 51], bidrectional MR (with the use of many MR methods) does not perform well due to the following reason: assuming the true causal direction is from *X* to *Y* for two traits *X* and *Y*, if an SNP is causal to *X* (and the sample sizes are large enough), the SNP is associated with both *X* and *Y*, and thus may be considered as a candidate IV for both traits; when the SNP is used as an IV for direction *X* to *Y*, it will confirm the causal association; however, if it is used as an IV for direction *Y* to *X*, it will also yield a non-zero estimate of the causal effect of *Y* on *X*, leading to an incorrect conclusion. A naive remedy is to remove any SNP associated with both traits, but it leads to not only loss of power (with fewer SNPs as IVs), but also biased inference (e.g., towards the causal direction *X* to *Y* if the truth is *Y* to *X* and if the GWAS sample size or power for *X* is much larger than for *Y*) [38]. Here we adopt a simple but effective screening/filtering rule based on a simple heuristic: no SNP will be used as an IV for both traits, because no SNP can be valid for both traits. If an SNP is associated with both traits, by Steiger’s method, we use it only for the trait with which its correlation is larger than that with the other trait (because it is more likely to be a valid IV for the chosen trait) [38]. Furthermore, there are some new MR methods, such as constrained maximum likelihood (MRcML) [52], that are more robust to both uncorrelated and correlated pleiotropy. Our second goal here is to show that, by incorporating MR-cML and the IV screening rule in reciprocal MR, the resulting method, still called MR-cML for simplicity, performs well, in fact almost identically to CD-cML if their modeling assumptions hold; otherwise, CD-cML controls type I error better and is more conservative.

With the two robust and powerful methods, we show their application to infer bi-directional relationships, which has been largely neglected in the literature. It is notable that inferring bidirectional causal relationships is far more challenging than uni-directional ones: for example, for the first time we point out that a bi-directional causal relationship generates a few new scenarios, in which either the InSIDE assumption or the plurality condition required by many existing robust MR methods will be violated (e.g. IVW (random effect), Egger regression [3] and RAPS [56] for the former; our cML methods, MR-ContMix [9], MR-Mix [36], MR-Lasso [7] and MR-Weighted Mode [19] for the latter). We applied the methods to 48 risk factor-complex disease pairs with 12 cardiometabolic risk factors, 3 cardiometabolic diseases (T2D, Stroke and CAD), and asthma (more as a negative control), identifying some interesting bidirectional causal relationships, such as between BMI and CAD, and between BMI and T2D [18].

## 2 Results

### 2.1 Overview of Methods

Given two traits, *X* and *Y*, and two independent GWAS datasets for the two traits, our goal is to infer their possibly *bi-directional* causal relationship. One of the most challenging issues is that we have a *hidden* (i.e. unobserved) confounder (or equivalently, an aggregate of many hidden confounders) denoted by *U*, which is associated with both *X* and *Y* with effect sizes *θ*_*UX*_ and *θ*_*UY*_ respectively. There are three possible sets of candidate SNPs to be used as IVs: (1) {*g*_*X*_} is the set of valid IVs for *X*, having direct effects *α*’s only on *X* ; (2) {*g*_*Y*_} is the set of valid IVs for *Y*, having direct effects *β* ’s only on *Y* ; (3) {*g*_*B*_} is the set of invalid IVs directly influencing both *X* and *Y*, and possibly *U*, with direct effects *γ*’, *η*’s, and *ξ* ’s respectively. Figure 1 illustrates a true causal model, in which *θ*_*XY*_ and *θ*_*YX*_ are the causal effects between the two traits, the unknown parameters of interest.

**Figure 1:**
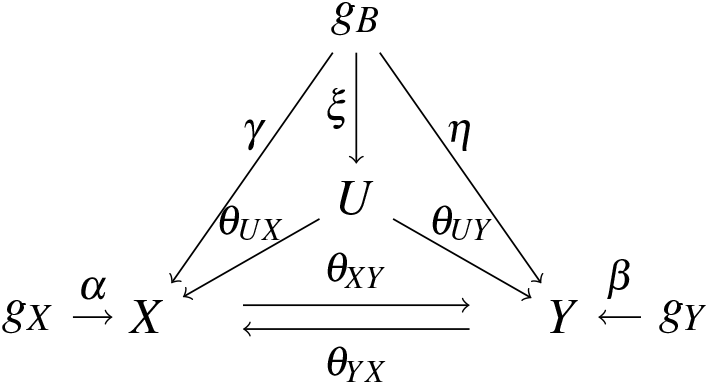
True causal model.

Define the (population) Pearson correlations between each candidate SNP/IV *g* and two traits as *ρ*_*Xg*_ = *corr*(*X, g*) and *ρ*_*Yg*_ = *corr*(*Y, g*). It is shown in Methods that

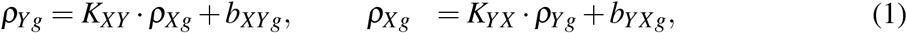

with 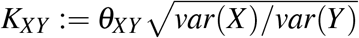 and 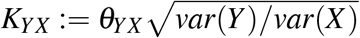.

If there exists causal direction from *X* to *Y*, we have | *K*_*XY*_ | < 1 (under a suitable condition) and *K*_*XY*_ ≠ 0, which can be used to infer the causal direction of *X* to *Y*. Similarly, we use | *K*_*YX*_ | < 1 and *K*_*YX*_ ≠ 0 to infer the causal direction of *Y* to *X*. The idea is similar to that used by other CD methods, i.e. Steiger’s method based on a single valid IV, CD-Ratio on multiple valid IVs and CD-Egger on multiple possibly invalid IVs without correlated pleiotropy (i.e. when the InSIDE assumption holds) [21, 51].

Since *K*_*XY*_ and *K*_*YX*_ are unknown, we propose a constrained maximum likelihood, called CD-cML, to infer the two parameters and thus the causal direction. Briefly, based on the given GWAS (summary) data, we calculate the sample (Pearson) correlations between each candidate SNP/IV and each trait, say *r*_*Yg*_ and *r*_*Xg*_, which are asymptotically normal and consistent for the (population) correlations *ρ*_*Yg*_ and *ρ*_*Xg*_; with eq (1), we can write down the normal-based log-likelihood under the constraint that the number of invalid IVs is equal to a given integer, say *m*_*I*_ ≥ 0. We try each possible value of *m*_*I*_, then consistently select the best one based on the Bayesian Information Criterion (BIC). The resulting constrained maximum likelihood estimates (cMLEs), say 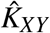 and 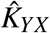, are consistent for *K*_*XY*_ and *K*_*YX*_ respectively, and are asymptotically normal. Hence we can construct a normal-based confidence interval for *K*_*XY*_ and *K*_*YX*_ respectively, thus drawing inference on the two possible causal directions from *X* to *Y* and from *Y* to *X*.

A similar method, called MR-cML, has been used to estimate *θ*_*XY*_ or *θ*_*YX*_ in the framework of MR [52]. It is noted that MR-cML performs well under correlated pleiotropy. Here we also propose applying MR-cML to reciprocal MR to infer both *θ*_*XY*_ and *θ*_*YX*_, and thus infer a possibly bi-directional causal relationship between *X* and *Y* ; for simplicity, we still call such a reciprocal MR as MR-cML.

It is noted that MR and CD methods are related but different: For example, for direction of *X* to *Y*, MR methods are based on inferring whether *θ*_*XY*_ = 0; in contrast, CD methods are based on whether *both K*_*XY*_ = 0 and | *K*_*XY*_ | < 1. Accordingly, because of the second constraint, we expect that sometimes CD-cML will be more conservative than MR-cML in terms of yielding smaller type I error and lower power.

We also propose a simple but yet effective method for SNP/IV screening based on a simple heuristic: none of SNPs can be a valid IV for both *X* and *Y*. Thus, if an SNP is (marginally) associated with both traits, we will use it only for the trait with which it is more correlated than with the other trait. This screening rule is just a simple application of Steiger’s method [21], and was mentioned in [38]. This screening rule is especially useful in the presence of bidirectional causal relationships. For example, when inferring whether there is a causal direction from *X* to *Y* in Figure 1, if *θ*_*YX*_ ≠ 0, all IVs in set {*g*_*Y*_} are associated with trait *X* and thus are candidate IVs, though they are all invalid IVs; the screening rule will eliminate them as IVs (because they will be more highly correlated with trait *Y* than with trait *X* ; see Methods). Now we consider what happens if they are indeed used as IVs. Assuming the set size |{*g*_*Y*_}| is larger than that of the valid IV set, |{*g*_*X*_}|, the invalid IV set {*g*_*Y*_}forms the largest (i.e. plurality) group in (incorrectly) estimating *θ*_*XY*_ as 1*/θ*_*YX*_ (asymptotically); in other words, they lead to the violation of the plurality condition required by cML (and several other MR methods, such as MR-ContMix [9], MR-Mix [36], MR-Lasso [7] and MR-Weighted Mode [19]). In addition, they will also lead to the violation of the InSIDE assumption: it is easy to verify that for any *g* ∈{*g*_*Y*_}, its effect size on trait *X* is *β*_*Xg*_ = *θ*_*YX*_ *β*_*Yg*_, which is clearly correlated with *β*_*Yg*_, its direct effect on *Y*.

There is another source leading to the violation of the InSIDE assumption, in addition to the more widely recognized one with *ξ* ≠ 0 (i.e. some IVs are correlated with the hidden confounder) and the one pointed above. It is due to bi-directional causal relationships, again demonstrating that it is more challenging to deal with bi-directional relationships than with unidirectional ones. Consider the causal direction of *X* to *Y* : even if *ξ* = 0 but if *θ*_*YX*_ ≠ 0, any SNP *g* ∈ {*g*_*B*_} would lead to the violation of InSIDE, because its association strength with *X* and its direct effect size on *Y* respectively would be *γ* + *ηθ*_*YX*_ and *η*, which are clearly correlated. The violation of the InSIDE assumption will lead to biased inference by several popular random-effects model-based methods (that treat direct effects as random effects), such as IVW (random effect), Egger regression [3] and RAPS [56].

Finally we apply the data perturbation (DP) scheme of [52] for better finite-sample inference: it accounts for uncertainty in selecting invalid IVs in CD- and MR-cML, leading to better control of type I errors. We suffix a method with “-DP” and “-S” to refer its use of data perturbation and IV screening respectively; by default, DP and IV screening are always used unless specified otherwise.

### 2.2 Simulations

We generated simulated data following the true causal model (1). There were 15, 10 and 10 SNPs/IVs in sets {*g*_*X*_ }, {*g*_*Y*_} and {*g*_*B*_}, respectively, with effect sizes ranging in (−0.3, −0.2) and (0.2, 0.3) (from the corresponding uniform distributions) respectively. For more correlated pleiotropy, we generated *ξ* ’s from a uniform distribution in the range of (−0.1, 0.1) or (−0.2, 0.2); otherwise, we set *ξ* ’s at 0. The MAFs of the SNPs were 0.3. The random errors *ε*_*X*_ and *ε*_*Y*_ were independently drawn from *N*(0, 1), and *ε*_*U*_ was from *N*(0, 2). We considered various combinations of the true causal effect sizes of *θ*_*XY*_ and *θ*_*YX*_ in the range from 0 to 0.3. For each set-up, we generated 500 pairs of two independent GWAS samples, one for each of the two traits and each of sample size *n* = 50000. For each dataset, we calculated the summary statistics for the two traits. We set the significance cutoff at 0.05/35 to select relevant IVs for both traits before applying any CD and MR methods.

We summarize the main results in terms of (empirical) type I error and power in Figures 2 and 3. The top-left panels (for “*θ*_*XY*_ = 0, X to Y”) show (empirical) type I error for the direction of *X*→ *Y* ; when *θ*_*YX*_ = 0 in the right panels, it shows type I error for the direction *Y* →*X* ; otherwise, it is for (empirical) power. In general, MR-cML and CD-cML (referring the version with data perturbation and IV screening by default, i.e. “-DP-S”, unless specified otherwise) performed almost identically: they could control type I error satisfactorily while having high power. In contrast, all other methods, namely CD-Ratio, CD-Egger and combining (single-SNP-based) Steiger’s method over multiple IVs (by majority voting, “-MV”), could not control type I error, and might have low power. Interestingly, CD-Egger had largely inflated type I error rates even when none of the IVs were correlated with the hidden confounder (i.e. *ξ* = 0) unless both *θ*_*XY*_ = *θ*_*YX*_ = 0 as shown in Figure 2. This might sound surprising, but convincingly showed that detecting bi-directional causal relationships is much more challenging than detecting uni-directional ones. As explained in Methods, even if *ξ* = 0, there were other two possible reasons why the InSIDE assumption required by Egger regression would be violated. For example, let’s consider causal direction from *X* to *Y* when *θ*_*XY*_ = 0 but *θ*_*YX*_ ≠ 0: any IV in {*g*_*Y*_}or {*g*_*B*_} would lead to the violation of the InSIDE assumption. Note that, although the proposed screening rule would eliminate using SNPs from {*g*_*Y*_}, some from {*g*_*B*_} would remain. On the other hand, when the SNPs from *g*_*B*_ were correlated with the hidden confounder (with *ξ* ≠ 0), the InSIDE assumption would always be violated, leading to inflated type I error of CD-Egger as shown in Figure 3. Notably, both MR- and CD-cML, equipped with the screening rule, did not suffer from any of these problems, partly because those parameters were treated as fixed nuisance parameters, instead of random effects in Egger regression.

**Figure 2:**
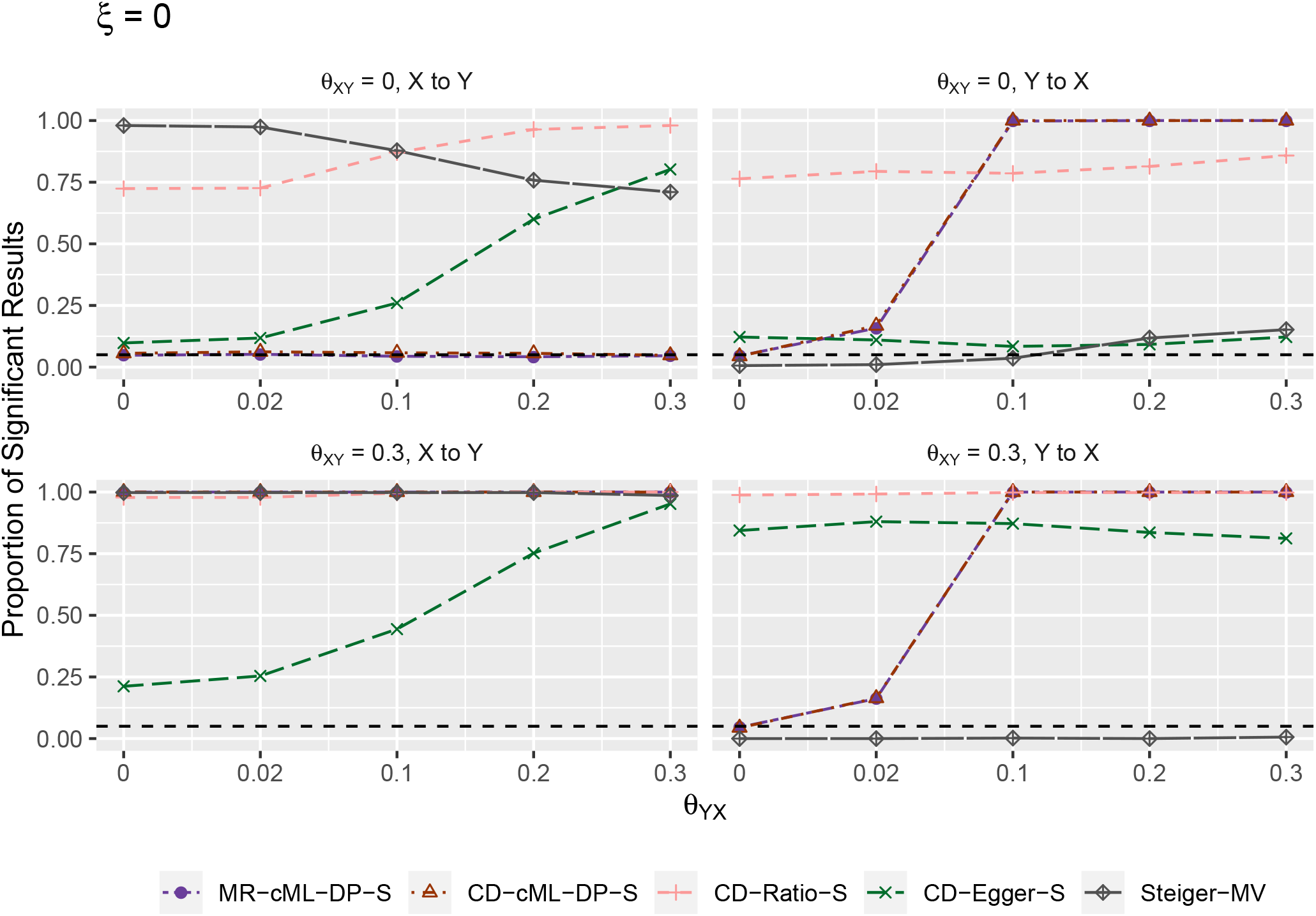
Empirical type-I error and power (y-axis) when *ξ* = 0 (i.e. no IV correlated with the hidden confounder), *θ*_*XY*_ = 0 (top panels) and *θ*_*XY*_ = 0.3 (bottom) for various values of *θ*_*YX*_ (x-axis). The left panels show results for the direction *X* → *Y*, the right ones show results for the direction *Y* → *X*.

**Figure 3:**
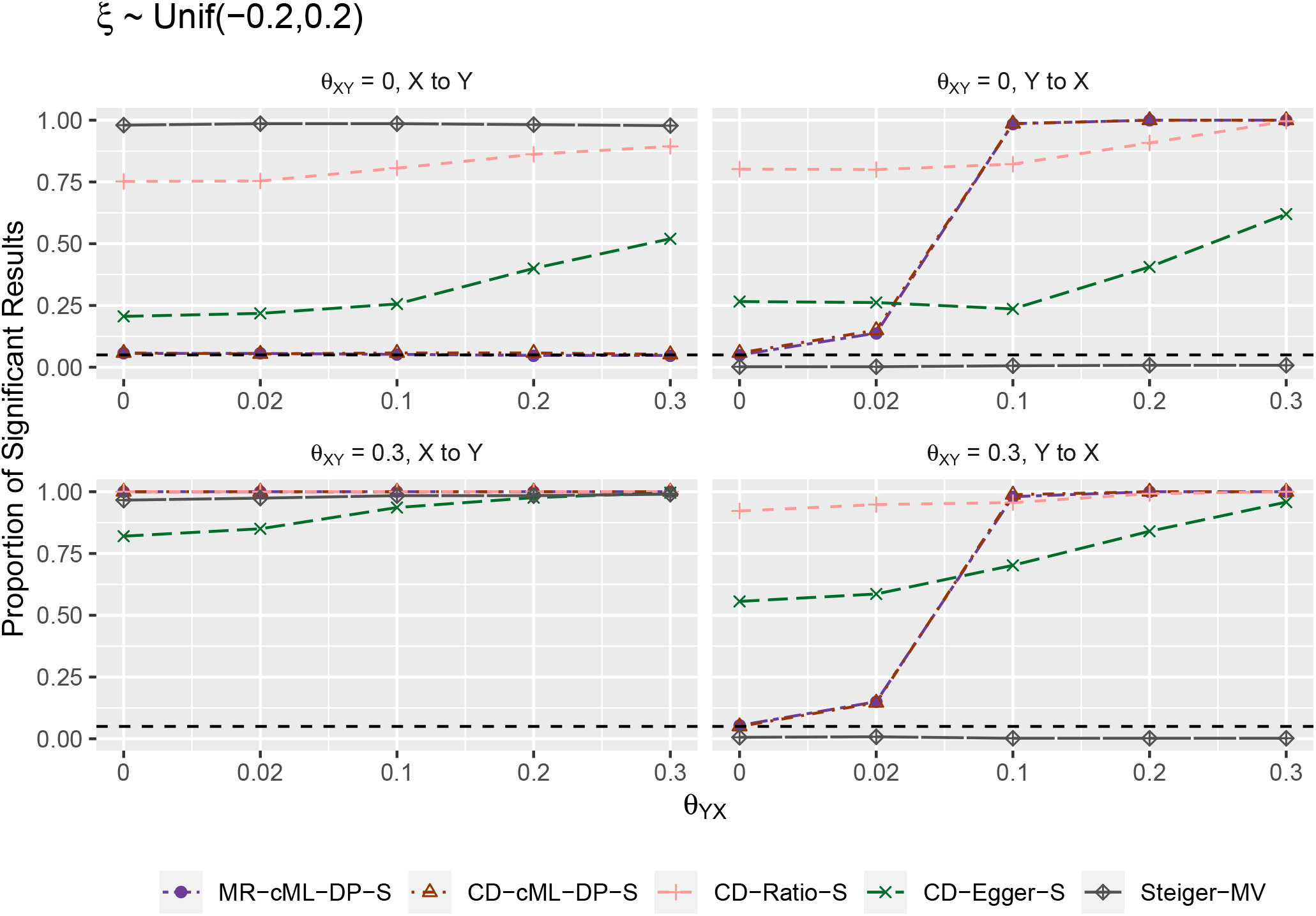
Empirical type-I error and power (y-axis) when *ξ* was from a uniform distribution (i.e. some IVs correlated with the hidden confounder, implying InSIDE is violated), *θ*_*XY*_ = 0 (top panels) and *θ*_*XY*_ = 0.3 (bottom) for various values of *θ*_*YX*_ (x-axis). The left panels show results for the direction *X* → *Y*, the right ones show results for the direction *Y* → *X*.

Figures 4-7 show the empirical distributions of the parameter estimates from the methods under various simulation set-ups. It is confirmed that equipped with the IV screening and data perturbation, both MR-cML and CD-cML always gave (almost) unbiased estimates of the causal parameter (*θ*) and the correlation ratio parameter *K*) respectively. In contrast, CDEgger was biased except when the InSIDE assumption held (i.e. *ξ* = 0 and no causal effect from the other direction), while CD-Ratio was in general biased in the presence of invalid IVs. Furthermore, CD-cML and MR-cML were much more efficient in yielding estimates with much smaller variances than those of CD-Ratio and CD-Egger.

**Figure 4:**
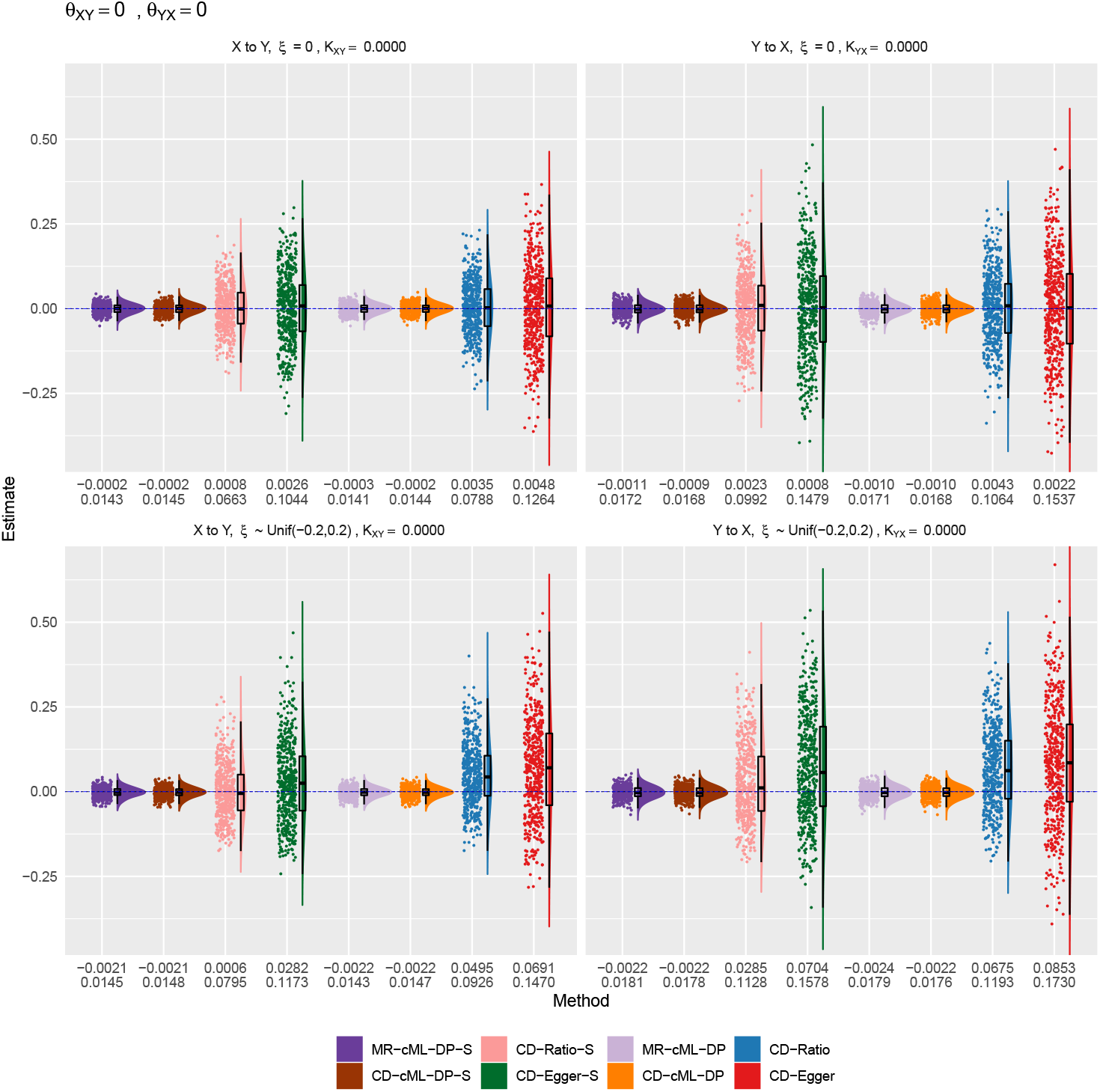
Empirical distributions of the estimates of the correlation ratio *K* for CD methods and the causal effect *θ* for MR method, when true *θ*_*XY*_ = 0 and *θ*_*YX*_ = 0. The top and bottom panels show results for *ξ* = 0 (i.e. no IV correlated with the hidden confounder) and *ξ* from a uniform distribution (i.e. some IVs correlated with the hidden confounder, implying InSIDE is violated), respectively. The left panels show the estimates for the causal direction of *X*→ *Y*, while the right ones show that for *Y* → *X*. The horizontal dashed lines are for the true values of *K* and *θ*. The two rows of the numbers under each panel give the sample mean and standard deviation of the estimates from each method.

**Figure 5:**
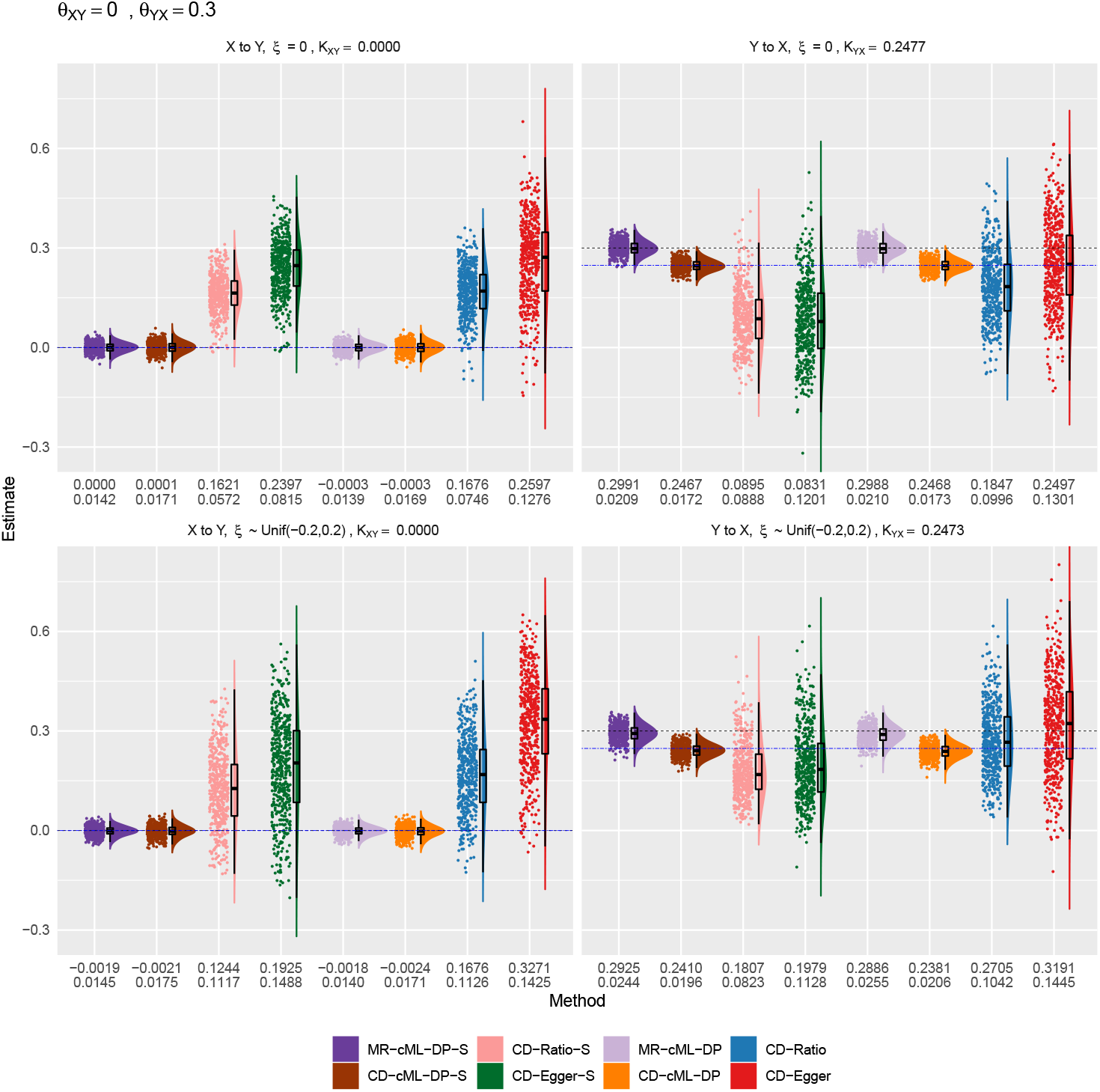
Empirical distributions of the estimates of the correlation ratio *K* for CD methods and the causal effect *θ* for MR method, when true *θ*_*XY*_ = 0 and *θ*_*YX*_ = 0.3. The top and bottom panels show results for *ξ* = 0 (i.e. no IV correlated with the hidden confounder) and *ξ* from a uniform distribution (i.e. some IVs correlated with the hidden confounder, implying InSIDE is violated), respectively. The left panels show the estimates for the causal direction of *X* → *Y*, while the right ones show that for *Y*→ *X*. The horizontal dashed lines are for the true values of *K* (in blue) and *θ* (in black). The two rows of the numbers under each panel give the sample mean and standard deviation of the estimates from each method.

**Figure 6:**
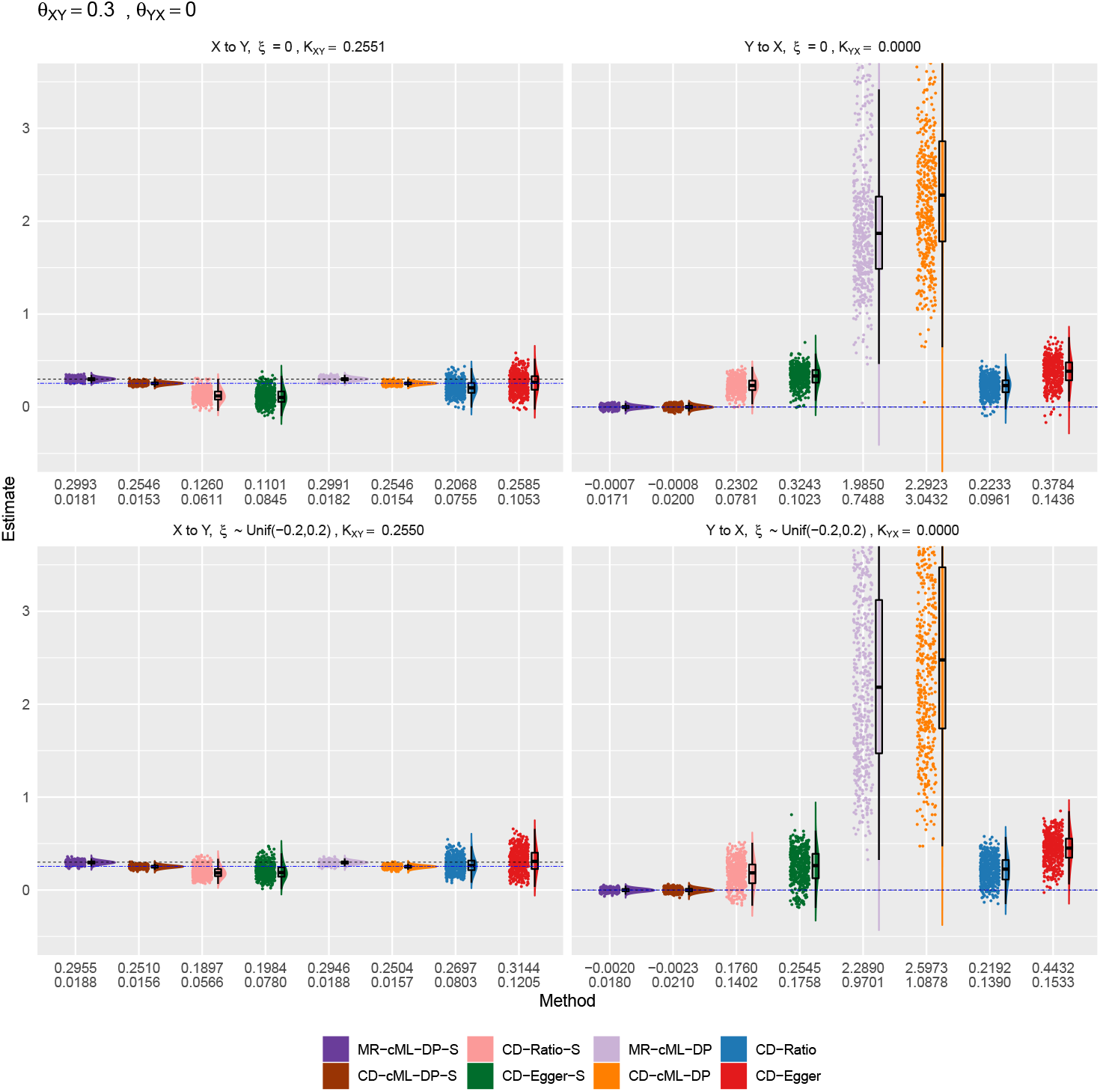
Empirical distributions of the estimates of the correlation ratio *K* for CD methods and the causal effect *θ* for MR method, when true *θ*_*XY*_ = 0.3 and *θ*_*YX*_ = 0. The top and bottom panels show results for *ξ* = 0 (i.e. no IV correlated with the hidden confounder) and *ξ* from a uniform distribution (i.e. some IVs correlated with the hidden confounder, implying InSIDE is violated), respectively. The left panels show the estimates for the causal direction of *X*→ *Y*, while the right ones show that for *Y*→ *X*. The horizontal dashed lines are for the true values of *K* (in blue) and *θ* (in black). The two rows of the numbers under each panel give the sample mean and standard deviation of the estimates from each method.

**Figure 7:**
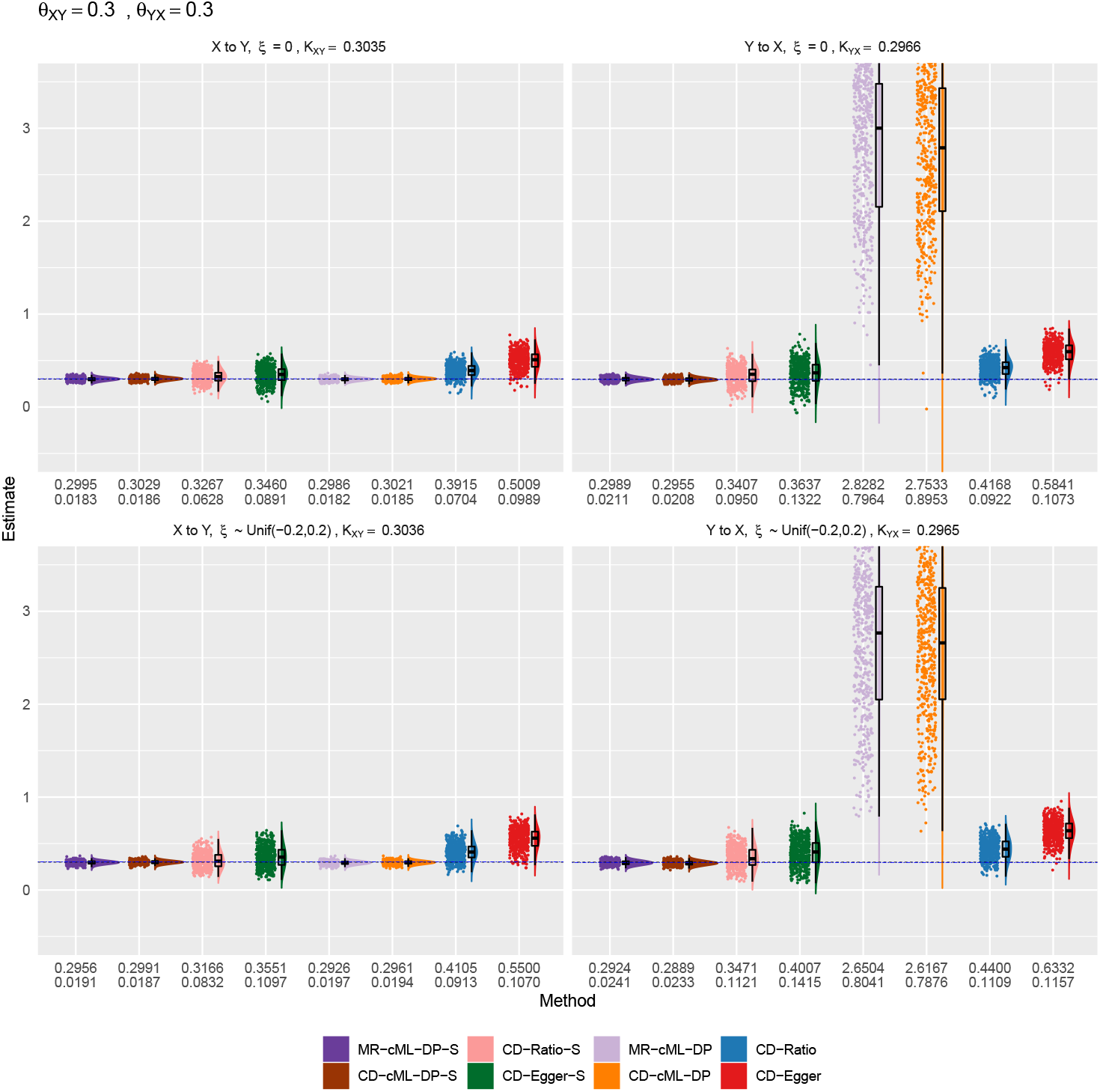
Empirical distributions of the estimates of the correlation ratio *K* for CD methods and the causal effect *θ* for MR method, when true *θ*_*XY*_ = 0.3 and *θ*_*YX*_ = 0.3. The top and bottom panels show results for *ξ* = 0 (i.e. no IV correlated with the hidden confounder) and *ξ* from a uniform distribution (i.e. some IVs correlated with the hidden confounder, implying InSIDE is violated), respectively. The left panels show the estimates for the causal direction of *X*→ *Y*, while the right ones show that for *Y*→*X*. The horizontal dashed lines are for the true values of *K* (in blue) and *θ* (in black). The two rows of the numbers under each panel give the sample mean and standard deviation of the estimates from each method.

We also illustrate the significant role of the IV screening rule: without it both MR- and CD- cML could be biased. As discussed in Methods and shown in Figures 6 and 7, when considering the direction of *Y* to *X* with *θ*_*XY*_ ≠ 0, without screening rule, the SNPs in {*g*_*X*_} would be used (as invalid IVs) and became the plurality group (due to | {*g*_*X*_ }| = 15 *>* 10 = | {*g*_*Y*_ }|, thus estimating *θ*_*YX*_ as 1*/θ*_*XY*_ incorrectly. Again this problem demonstrates an additional challenge with bi-directional relationships. It is also noted that, in these situations, the estimated *K*’s were much larger than 1, leading to much smaller type I error (and lower power) of CD-cML than MR-cML.

More results for the methods with or without data perturbation and/or IV screening are provided in the Supplementary.

### 2.3 Real Data Examples

We studied possibly bi-directional causal relationships between 12 cardiometabolic risk factors and 4 common diseases, and between pairs of the 4 common diseases, including 3 cardiometabolic diseases, namely coronary artery disease (CAD) [43], Stroke [30], type 2 diabetes (T2D) [31], and asthma (largely serving as a negative control) [15]. The 12 risk factors are triglycerides (TG), low-density lipoprotein cholesterol (LDL), hight-density lipoprotein cholesterol (HDL) [48], Height [49], body-mass index (BMI) [28], body fat (BF) [29], birth weight (BW) [23], diastolic blood pressure (DBP), systolic blood pressure (SBP) [17], fasting glucose (FG) [16], Smoke and Alcohol [27]. Based on the existing literature, [32] partitioned the 48 risk factor-disease pairs into four groups, representing likely “causal”, “correlated”, “unrelated” and “non-causal” relationships.

We used R package `TwoSampleMR` to extract and pre-process the GWAS summary statistics. For each pair of traits *X* and *Y*, at the p-value cutoff 5 ×10^−8^, we first extracted all SNPs significant with *X*, denoted by *I*_*X*_, and all SNPs significant with *Y*, denoted by *I*_*Y*_. Let *I*_*U*_ = *I*_*X*_ ∪ *I*_*Y*_ be the combined set of the significant SNPs for at least one of the two traits, then we extracted the summary statistics for all SNPs in *I*_*U*_. For each SNP in *I*_*U*_, we defined its combined p-value across the two traits as *p*_*c*_ = *p*_*X*_ · *p*_*Y*_, where *p*_*X*_ and *p*_*Y*_ were the p-values for the two traits. We applied function `clump data` for clumping, using its default setting with distance 10000kb, *r*^2^ = 0.001, the European population as the reference panel, and *p*_*c*_’s as p-values for the SNPs. After clumping, we obtained a set of approximately independent SNPs, for which we had their GWAS summary statistics 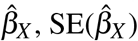 and 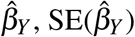.

We input the summary statistics into each CD or MR method, drawing conclusions according to the decision rules (see Methods). For cML, we used 10 random starting points to find cMLEs. For data perturbation, we set the number of perturbations *T* = 100. For all methods except Steiger’s method, for both directions, we used the Bonferroni-corrected significance level 0.05/48 ≈ 0.001 to construct confidence intervals. For Steiger’s method we show the results as (Proportion, Majority Vote). For example, for TG to CAD 50.5% SNPs were significant, for CAD to TG 18.4% SNPs were significant, and the rest 31.1% SNPs were not significant for either direction; Steiger-MV would conclude that TG to CAD as TRUE, and CAD to TG as FALSE. See Supplementary for detailed results.

As shown in Figure 8, many of the well-accepted causal relationships from a risk factor to a disease were confirmed by most of the methods, such as from TG to CAD, and LDL to CAD. There were also some interesting findings about causal effects from diseases to risk factors, for example, T2D to BMI, and T2D to FG identified by MR-cML, CD-cML and CD-Ratio, and Stroke to DBP by MR-cML and CD-cML. Between MR-cML and CD-cML, they gave mostly consistent results except for three pairs: 1) CD-cML, but not MR-cML, detected CAD to BMI. As shown in the Supplementary, the two methods gave the CIs of *K* and *θ* as (−0.199, −0.004) and (−0.081, 0.001) respectively, so the latter just barely covered 0. 2) MR-cML, but not CD- cML, identified two well-accepted causal relationships from BMI and FG to T2D. In both cases, the two methods gave the relatively wider CIs with that of CD-cML covering *K* = 1, indicating perhaps its lack of power and/or its being more conservative as discussed earlier. Finally, both CD-Ratio and CD-Egger, or only CD-Ratio, suggested a few more causal relationships from a disease to a risk factor, which however need to be further explored given the strong assumption of CD-Ratio with all valid IVs and of CD-Egger’s InSIDE assumption.

**Figure 8:**
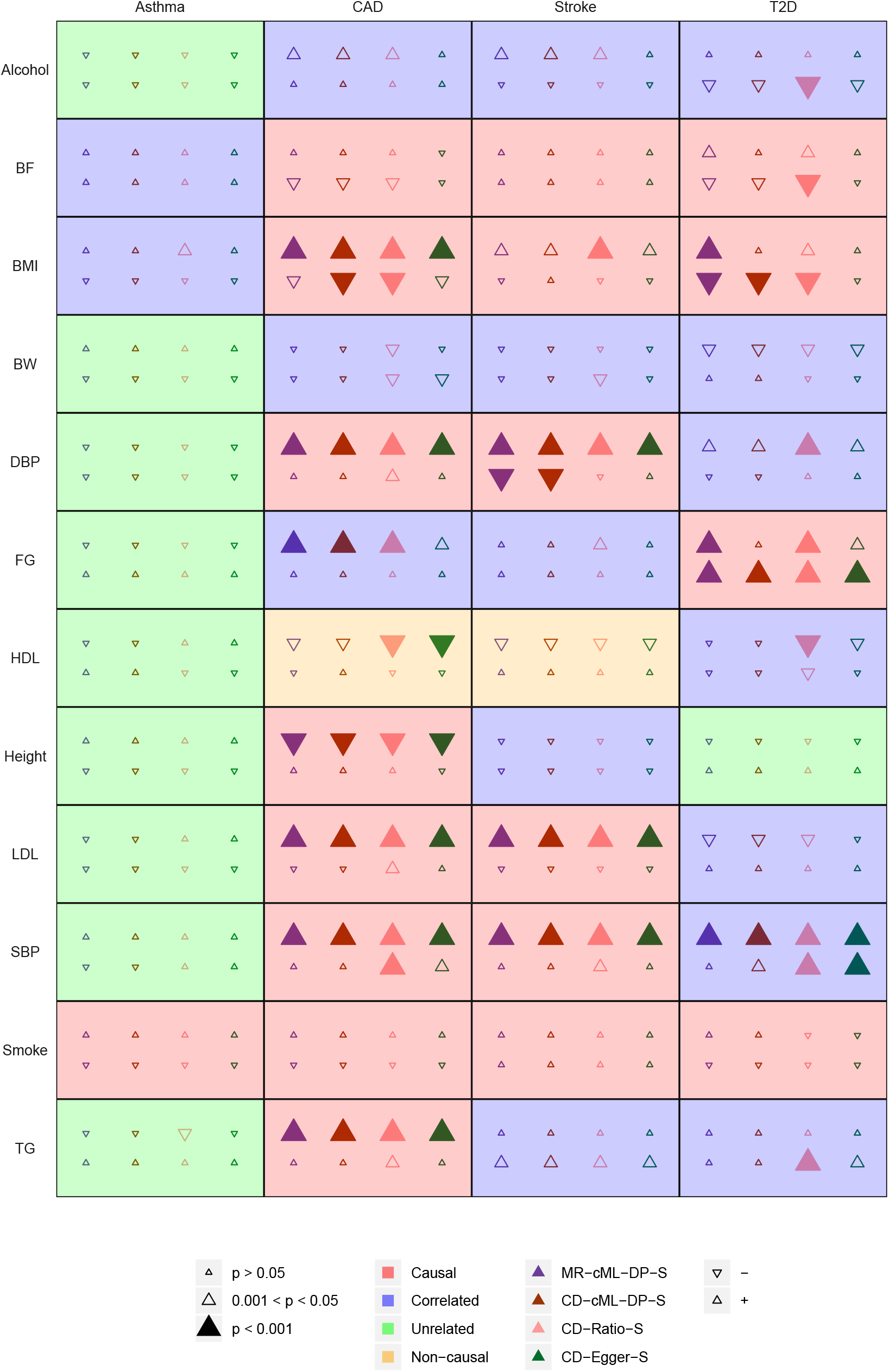
Causal relationship of 48 exposure-outcome pairs. As shown in the legends, the size, color and direction of a triangle represent the statistical significance, method being used, and the direction/sign of the estimated effect. In each cell, the first row is for the causal direction from risk factor to disease, and the second row is for the reverse.

In the Supplementary we show the detailed results between any two of the four diseases. Based on the Bonferroni-adjusted significance cutoff of 0.05/12 ≈ 0.004, all four methods identified three causal relationships: from CAD to Stroke, T2D to CAD and T2D to Stroke [18]. On the other hand, only CD-Ratio and CD-Egger suggested a reverse causation from Stroke to CAD. Finally, we note that MR-cML and CD-cML yielded consistent results.

## 3 Discussion

Inference for bi-directional causal relationships is far more challenging than for uni-directional ones, which has been the focus in the literature. In particular, in most MR analyses, a unidirectional causal relationship is assumed to be known and thus pursued. There are a few exceptions, though many considered bi-directional ones under the unrealistic assumption of having only valid IVs [40, 51]. The new work of [11] aimed for the same problem as considered here. However, there are some major differences. Foremost, although the two true causal models are similar, the technical approaches are substantially different: they estimate all the parameters related to various direct and indirect effect sizes among IVs, the hidden confounder and the two traits in the model, requiring much stronger modeling assumptions, such as the normality assumption on the distribution of the effect sizes, leading to not only a more complex estimation procedure (with a Bayesian approach), but also some non-identifiability for parameter estimation. In contrast, except for the few key parameters of interest, such as the causal effect sizes, we treat the majority of the other parameters as nuisance ones that are combined into two intercept parameters in our model (1), leading to a much simpler estimation procedure without any distributional assumption on these nuisance parameters. We also note that the major assumption in our cML method is the “plurailty condition” (that the valid IVs form the largest/plurality group in estimating the same causal parameter as their estimand), which is relatively weak because it does not even require the majority of the IVs to be valid. Nonetheless, for bi-directional causal relationships, it will always be violated for one trait if no screening is applied: since the valid IVs for one trait are invalid for the other trait, we have either an equal or even a larger number of invalid IVs to or than the number of valid IVs; the former set of invalid IVs correspond to the same (incorrect) causal estimand, implying that the set of valid IVs will not form the largest group in estimating the same (correct) causal estimand; that is, the plurality condition will be violated. In addition, this problem will be catastrophic to other methods based on modeling direct effects of invalid IVs as random effects, such as Egger regression, IVW-RE and RAPS, because it leads to the violation of the InSIDE assumption required by these methods. The simple screening rule based on the Steiger’s method is surprisingly effective in eliminate (or alleviate) such a problem.

The recent popularity of Steiger’s method may suggest that correlation-based CD methods should be more effective than MR in determining causal directions. Surprisingly, as shown here, equipped with the same screening rule and the same robust cML estimation method, we do not see better performance of CD-cML over MR-cML when their modeling assumptions, mainly of the plurality condition, hold. A possible explanation is the following: CD methods exploit the key relationship characterizing the correlations between each trait and each SNP/IV as defined by the *K* parameter, which however is derived from the same causal model used by MR. In fact, since *K* is proportional to the causal parameter *θ*, according to the invariance property and the asymptotic optimality of the maximum likelihood estimator, there should be no advantage in estimating either one over the other, thus their large-sample inference and the corresponding conclusions should be (asymptotically) equivalent if their modeling assumptions hold. However, since CD methods exploit the key condition | *K*| < 1, they are less likely to make type I errors (albeit with lower power) than MR when their modeling assumptions are violated, as shown in our simulations (Figures 6 and 7). Therefore, as evidenced in our real data application, given no guarantee of the validity of all modeling assumptions, including the plurality condition, for any given problem, CD-cML can serve as a more conservative alternative to MR-cML.

There are a few limitations with the current study. First, we have only considered the two-sample design with two independent GWAS datasets for the two traits. It will be useful to extend the methods to the case with overlapping individuals in the two GWAS datasets, or to the one-sample design with only one GWAS dataset for both the traits [32, 11]. Second, in our real data examples the SNPs/IVs were selected from the same two GWAS datasets for the two traits, which might introduce selection bias. Alternatively, we can use an independent GWAS dataset for SNP selection for each trait [55], or analytically account for selection effects [53, 25, 46]. Third, as usual in MR we only considered linear models, while non-linear modeling may be more flexible and gain power [5, 54]. Finally, and most importantly, any analysis method comes with its modeling assumptions, especially in causal inference with observational data. Although we feel that our modeling assumptions for our proposed CD- and MR-cML, mainly the plurality condition, are relatively weak, they may or may not hold in practice. It would be useful to develop model checking techniques, or apply alternative methods under different modeling assumptions for triangulation [26]. Only more real data applications can shed light on how likely these modeling assumptions, especially the plurality condition, would hold in practice.

## 4 Methods

### 4.1 Model

The true causal model depicted in Figure 1 can be expressed as

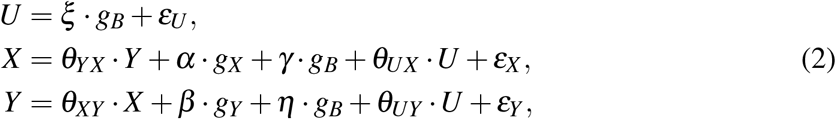

where the random errors *ε*_*U*_, *ε*_*X*_ and *ε*_*Y*_ are independent with each other. The reduced form of the true model is

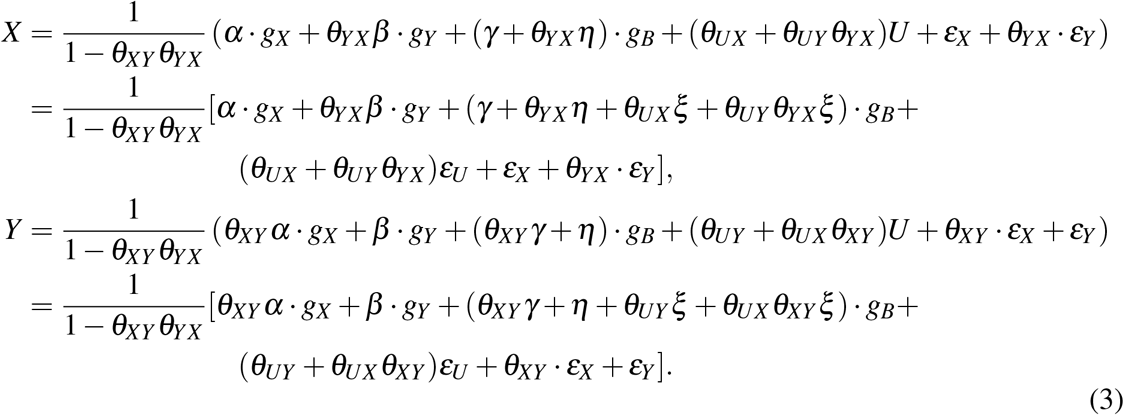

First, for an IV *g* in {*g*_*X*_ }, its (population) correlations with *X* and *Y* are

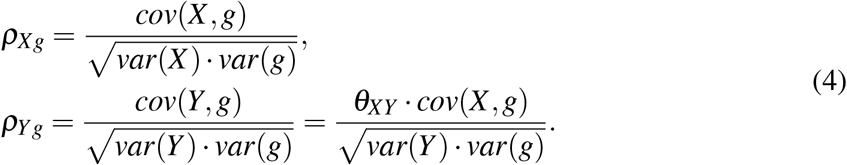

We have

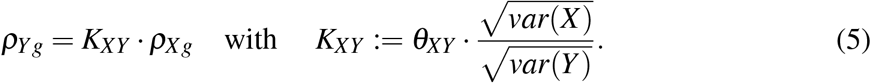

Under the key assumption 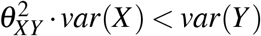, we have |*K*_*XY*_ | < 1. More discussions on why this assumption is often reasonable and how to empirically check this assumption are given in [51].

Second, for an IV *g* in {*g*_*Y*_ }, its correlations with *X* and *Y* are

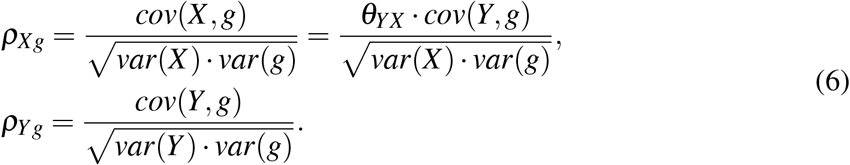

We have

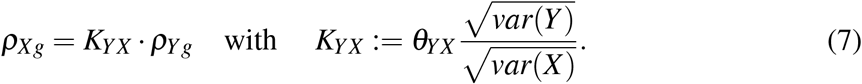

Again under the key assumption 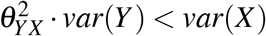, we have |*K*_*YX*_ | < 1.

Third, for an IV *g* in {*g*_*B*_}, its correlations with *X* and *Y* are

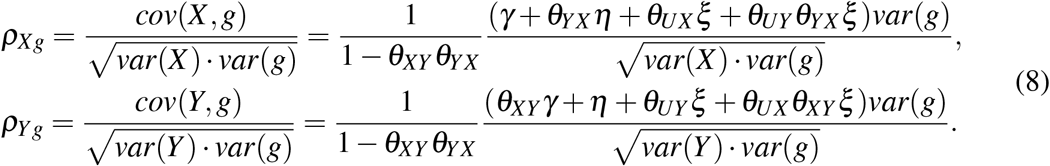

We have

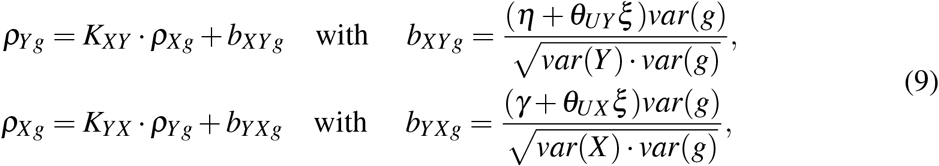

where *K*_*XY*_ and *K*_*YX*_ are defined as before, and satisfying |*K*_*XY*_| < 1 and |*K*_*YX*_ |< 1 under each of the two key assumptions respectively.

Note that *ξ* ≠ 0 induces correlations between *ρ*_*Xg*_ and *b*_*XYg*_, and between *ρ*_*Yg*_ and *b*_*YXg*_, through the shared term *ξ*, leading to the violation of the InSIDE assumption, as expected. However, even if *ξ* = 0, that *θ*_*YX*_ ≠ 0 or *θ*_*XY*_ ≠ 0 would still induce a correlation between *ρ*_*Xg*_ and *b*_*XYg*_ (through the shared term *η*), or between *ρ*_*Yg*_ and *b*_*YXg*_ (through shared *γ*), respectively, again leading to the violation of InSIDE. Furthermore, for *g* ∈ {*g*_*Y*_ }, if *θ*_*YX*_ ≠ 0, from equation (7) we have

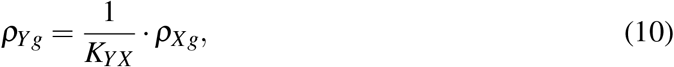

which means that if *g* ∈ {*g*_*Y*_ }is used as an IV to infer the causal direction from *X* to *Y*, it would incorrectly estimate *K*_*XY*_ as 1*/K*_*YX*_. In particular, if the set size |{*g*_*Y*_ }| is larger than that of the valid IV set, | {*g*_*X*_} |, it implies that the plurality condition will be violated when the SNPs in {*g*_*Y*_} are used as candidate IVs; this problem, completely due to considering a bi-directional relationship (i.e. when considering direction *X* to *Y* while allowing a causal direction of *Y* to *X*), can be avoided or alleviated by applying the IV screening rule to be discussed later. In summary, these various scenarios showcase unique challenges with analysis of bi-directional relationships.

Note that eq (10) can be rewritten as

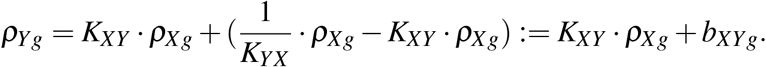

Hence, combining all possible cases for *g* ∈ {*g*_*X*_} ∪ {*g*_*Y*_} ∪ {*g*_*B*_}, eq (9) holds and can serve as a general model for statistical estimation.

Finally, it is noted that *K*_*XY*_ ≠ 0 (or *K*_*YX*_ ≠ 0) if and only if *θ*_*XY*_ ≠ 0 (or *θ*_*YX*_ ≠ 0). Accordingly we can infer causal directions based on whether *K*_*XY*_ and *K*_*YX*_ are non-zero and whether their absolute values are less than 1. Similarly, based on whether *θ*_*XY*_ and *θ*_*YX*_ are 0 or not, we can infer causal directions. Since these are unknown parameters, next we propose how to estimate them.

### 4.2 Constrained Maximum Likelihood

From a GWAS summary dataset for trait *X*, for each *g* in {*g*_*X*_ } ∪ {*g*_*Y*_ } ∪ {*g*_*B*_}, we have its estimated effect size 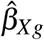 with standard error SE 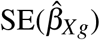 ;as discussed in [51], we can calculate the sample correlation *r*_*Xg*_ and its standard error SE(*r*_*Xg*_); similarly, for trait *Y*, we obtain its sample correlation *r*_*Yg*_ with standard error SE(*r*_*Yg*_).

For causal direction *X*→ *Y*, based on the consistency and asymptotic normality of the sample correlations *r*’s and their standard errors SE(*r*)’s, and general model (9), we write down the log-likelihood as

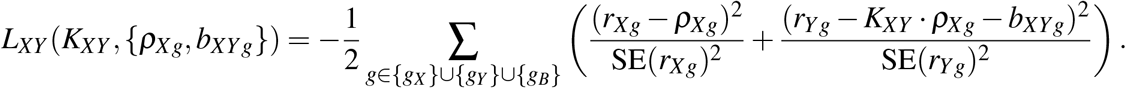

We apply the constrained maximum likelihood method [52]:

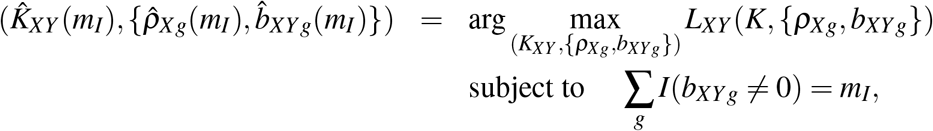

where *m*_*I*_ is a specified number of invalid IVs. Since *m*_*I*_ is unknown, we try *m*_*I*_ ∈ *ℳ* = {0, 1, …, *m−* 2 }, where *m* is the total number of the IVs being used, then use the Bayesian Information Criterion (BIC) to select the best *m*_*I*_. The BIC with *m*_*I*_ is

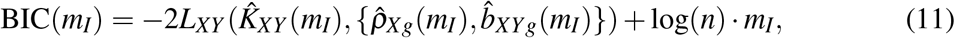

where *n* can be an any integer between *N*_1_ and *N*_2_, the sample sizes for the two GWAS for *X* and *Y* respectively, though we recommend using *n* = min(*N*_1_, *N*_2_). We select 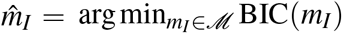, and estimate the set of invalid IVs as 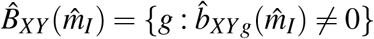.

Under the plurality condition (i.e. that the valid IVs form the largest group giving the same (asymptotic) estimate of *K*_*XY*_) and that the two GWAS sample sizes are comparable, as in [52], we can consistently select *m*_*I*_ (as the true number of invalid IVs), and the resulting constrained maximum likelihood estimate (cMLE) 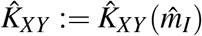 is consistent for the true value of *K*_*XY*_ and (asymptotically) normally distributed, as shown below.

#### Assumption 1.

*(Plurality condition*.*) Suppose that* 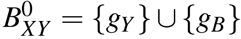*is the index set of the true invalid IVs for direction X* → *Y, with* 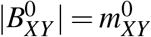. *For any B* ⊆ {1, …, *m*} *and* 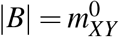, *If* 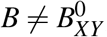, *then the* 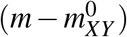 *ratios* {*b*_*XYg*_*/ρ*_*Xg*_, *g* ∈ *B*^*c*^} *are not all equal*.

#### Assumption 2.

*(Orders of the variances and sample sizes*.*) There exist positive constants l*_*X*_, *l*_*Y*_, *l*_*N*_ *and u*_*X*_, *u*_*Y*_, *u*_*N*_ *such that we have l*_*X*_ */N*_1_ ≤ *SE*(*r*_*Xg*_)^2^ ≤ *u*_*X*_ */N*_1_, *l*_*Y*_ */N*_2_ ≤ *SE*(*r*_*Yg*_)^2^ ≤ *u*_*Y*_ */N*_2_, *and l*_*N*_ · *N*_2_ ≤ *N*_1_ ≤ *u*_*N*_ · *N*_2_ *for g* = 1, …, *m*.

#### Theorem 1.

*Under Assumptions 1 and 2*, 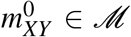, *we have* 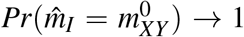 *and* 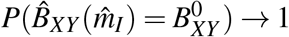. *Furthermore, the* 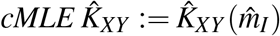 *is consistent and asymptotically normal:*

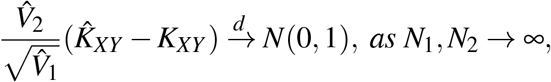

*Where*

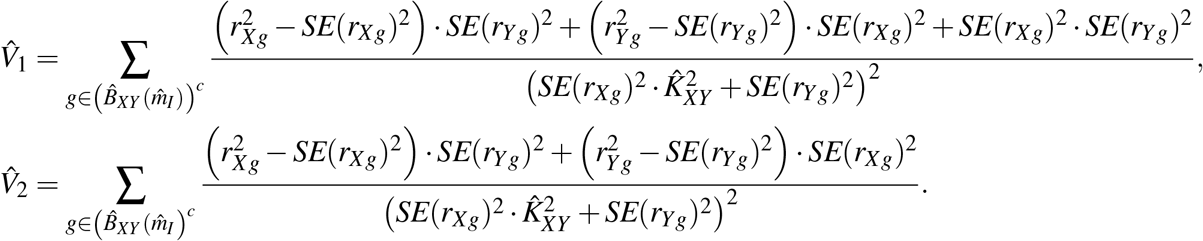

The proof for the selection consistency of BIC is similar to that for MR-cML [52], while that for the consistency and asymptotic normality of the cMLE parallels the proof of Theorem 3.3 in [56], for which the conditions are satisfied, including that here we consider only a fixed/finite *m*. We note that we can also use the information matrix to consistently estimate the variance of 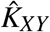, and thus construct a normal confidence interval (CI) for *K*_*XY*_ (or calculate a p-value). We can similarly estimate *K*_*YX*_ and draw inference.

As in [52], first, we can similarly apply a fast iterative algorithm to obtain the cMLE. Second, we propose using model averaging, instead of model selection for a single best *m*_*I*_, to account for model selection uncertainty and achieve better finite-sample performance. Third, we apply data perturbation to further improve performance for finite-sample inference, especially to better control type I error.

We note that our proposed the cML method, called CD-cML here, is similar to MR-cML in [52]. The main difference is that in MR-cML, the effect size estimates 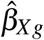 and 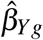 replace the sample correlations to estimate the causal effect *θ*_*XY*_ or *θ*_*YX*_, instead of *K*_*XY*_ or *K*_*YX*_ ; all other aspects remain the same.

### 4.3 IV Screening

For direction *X* → *Y* we identify the initial set of the significant SNPs associated with *X*, denoted by *I*_*X*_ ; for direction *Y* → *X* we identify the initial set of the significant SNPs with *Y*, denoted by *I*_*Y*_. Then for each SNP in the intersection, *g* ∈ *I*_*X*_ ∩ *I*_*Y*_, if |*r*_*Xg*_| ≥ |*r*_*Yg*_| we keep SNP *g* in *I*_*X*_ and remove it from *I*_*Y*_ ; if |*r*_*Xg*_| < |*r*_*Yg*_| we keep it in *I*_*Y*_ and remove it from *I*_*X*_.

### 4.4 Other Methods and Decision Rules

We will compare the proposed CD-cML with MR-cML and other CD methods: Steiger’s method, CD-Ratio (like MR-IVW) and CD-Egger (like MR-Egger) [51]. Since Steiger’s method is based on a single SNP/IV, we propose a majority voting (MV) method to combine its results across multiple SNPs/IVs. Each IV would conclude with one of three possible results: (1) no causal relationship between *X* and *Y* ; (2) *X* has a causal effect on *Y* ; (3) *Y* has a causal effect on *X*. We would go with the conclusion reached by the majority of the IVs (and randomly break the ties if any), called Steiger-MV. It is clear that Steiger-MV cannot detect bi-directional relationships. In the Supplementary, we show the proportions of the IVs in the three conclusion groups respectively, called Steiger-Prop.

For any MR method, we first estimate 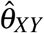 and SE 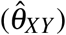 for *X Y*, and get a p-value *p*_*XY*_ ; for a given significance cutoff *α*, if *p*_*XY*_ < *α* we conclude *X* has a causal effect on *Y*, otherwise we do not. Similar we make a conclusion on whether there is a causal relationship of *Y* → *X*.

For any CD method except Steiger’s, for direction *X* → *Y*, we estimate 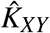 and 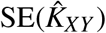, for *X* → *Y*, then for a given significant cutoff *α* then we construct a (1 − *α*) confidence interval of *K*_*XY*_ as 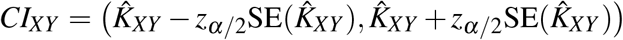, here *z*_*α*/2_ is the upper *α*/2 quantile of the standard normal distribution. If *CI*_*XY*_ is completely within (−1,0) or (0,1), we conclude that *X* has a causal effect on *Y*, otherwise *X* does not. Similarly, we make a conclusion about *Y* → *X*.

### 4.5 Simulation Setups

We independently generated 15 IVs in {g_*X*_} with *α*_1_, …, *α*_15_ from a uniform distribution Unif((−0.3, 0.2) ∪ (0.2, 0.3)); 10 IVs in {*g*_*Y*_} with *β*_1_, …, *β*_10_ from Unif((−0.3, 0.2) ∪ (0.2, 0.3)); and 10 IVs in {*g*_*B*_} with *γ*_1_, …, *γ*_10_ from Unif((−0.3, 0.2) ∪ (0.2, 0.3)), and *η*_1_, …, *η*_10_ from Unif((−0.3, 0.2) ∪ (0.2, 0.3)). We generated *ξ* ’s in two ways: i) set *ξ* ’s = 0 for no correlations with the confounder; ii) generated *ξ* ’s from Unif(− 0.1, 0.1) or Unif(− 0.2, 0.2) for correlations with the confounder.

The MAFs of the SNPs/IVs were all set as 0.3. *ε*_*X*_, *ε*_*Y*_ were independently drawn from N(0, 1), and *ε*_*U*_ from N(0,2). We tried different combinations of (*θ*_*XY*_, *θ*_*YX*_) = 0, {0.02, 0.1, 0.2, 0.3}× {0, 0.02, 0.1, 0.2, 0.3 }.

We generated two independent samples each of size *n* = *N*_1_ = *N*_2_ = 50000 from the reduced form (3) of the causal models for the two traits. Then we calculated and used subsequently the summary statistics for the two traits *X* and *Y* respectively. We set the significance cutoff at 0.05/35 to select relevant SNPs/IVs for both directions, and applied all methods for comparison.

## Supporting information

Supplementary Results

## Data and Code Availability

GWAS summary datasets being used in our real data examples are all publicly available in their corresponding references. Our proposed methods are implemented in R package `BiDirectCausal` which is available on GitHub at https://github.com/xue-hr/BiDirectCausal. Other R packages being used in this study are available with links in the Web Resources section.

## Supplementary Materials

In a Supplementary File we provide more detailed numerical results.

## Acknowledgements

This work was supported by NIH grants R01AG065636, R01 AG069895, RF1 AG067924, R01HL116720, R01GM113250 and R01GM126002, and by the Minnesota Supercomputing Institute at the University of Minnesota.

## Web Resources

TwoSampleMR, https://github.com/MRCIEU/TwoSampleMR

MRcML, https://github.com/xue-hr/MRcML

MRCD, https://github.com/xue-hr/MRCD

