## Supplementary Results for "Robust Inference of Bi-Directional Causal Relationships in Presence of Correlated Pleiotropy with GWAS Summary Data"

#### Contents

**S1 Full Simulation Results** **2**

**S2 Full Real Data Results** **17**
S2.1 48 Exposure-Outcome Pairs . . . . . . . . . . . . . . . . . . . . . . . . . . . . 17
S2.2 Pairs of 4 Diseases . . . . . . . . . . . . . . . . . . . . . . . . . . . . . . . 25

#### S1 Full Simulation Results

Figure S1: When $\theta_{XY} = 0, \xi = 0$, proportions of significant simulation results obtained by each methods for direction $X \to Y$ (left column) and $Y \to X$ (right column). The first row shows results for four main methods: MR-cML-DP-S, CD-cML-DP-S, CD-Ratio-S, and CD-Egger-S; the second row shows results for four methods without screening: MR-cML-DP, CD-cML-DP, CD-Ratio, and CD-Egger; the third row shows results for other five methods.

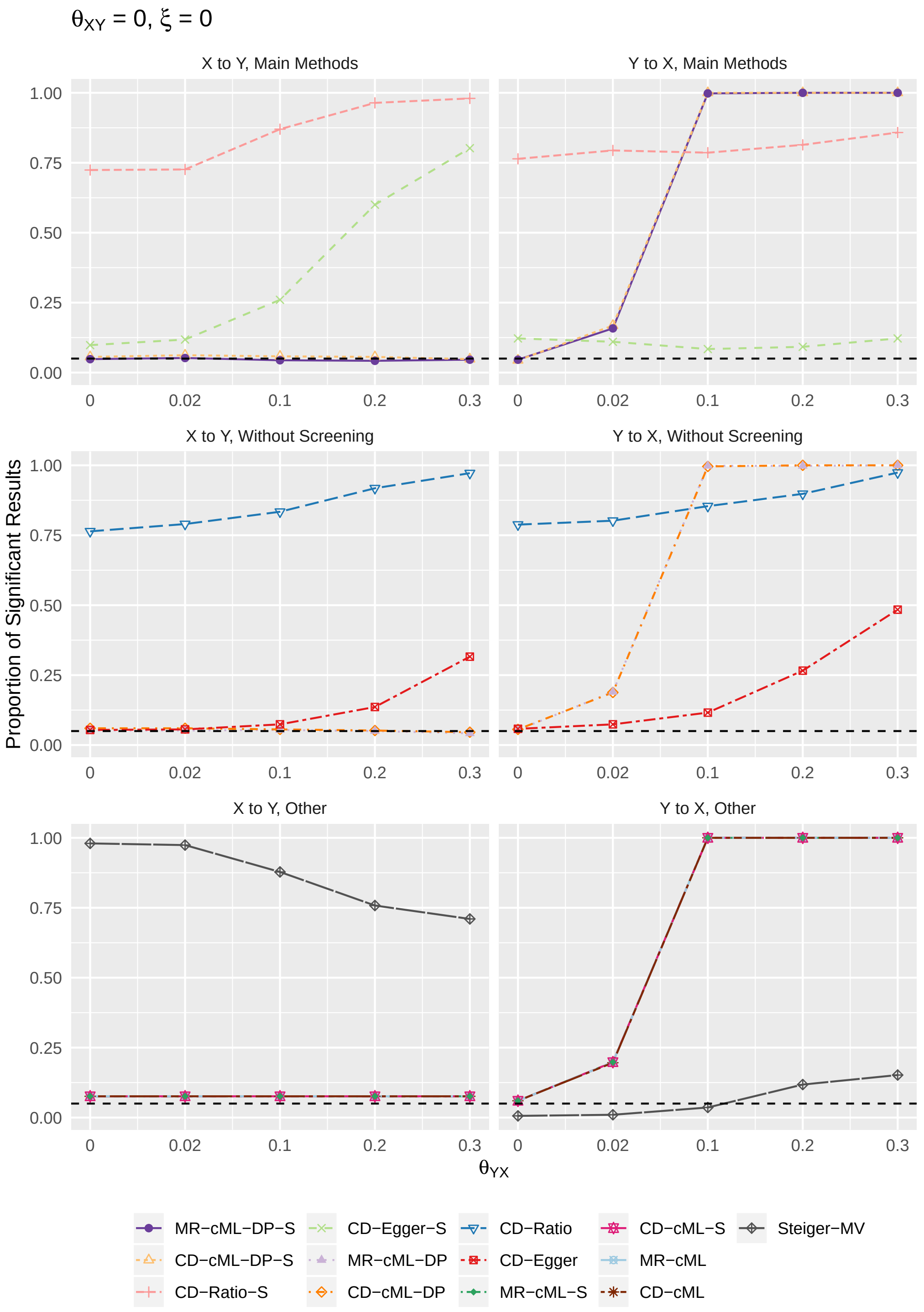

Figure S2: When $\theta_{XY} = 0, \xi \sim$ Unif(-0.1,0.1), proportions of significant simulation results obtained by each methods for direction $X \to Y$ (left column) and $Y \to X$ (right column). The first row shows results for four main methods: MR-cML-DP-S, CD-cML-DP-S, CD-Ratio-S, and CD-Egger-S; the second row shows results for four methods without screening: MR-cML-DP, CD-cML-DP, CD-Ratio, and CD-Egger; the third row shows results for other five methods.

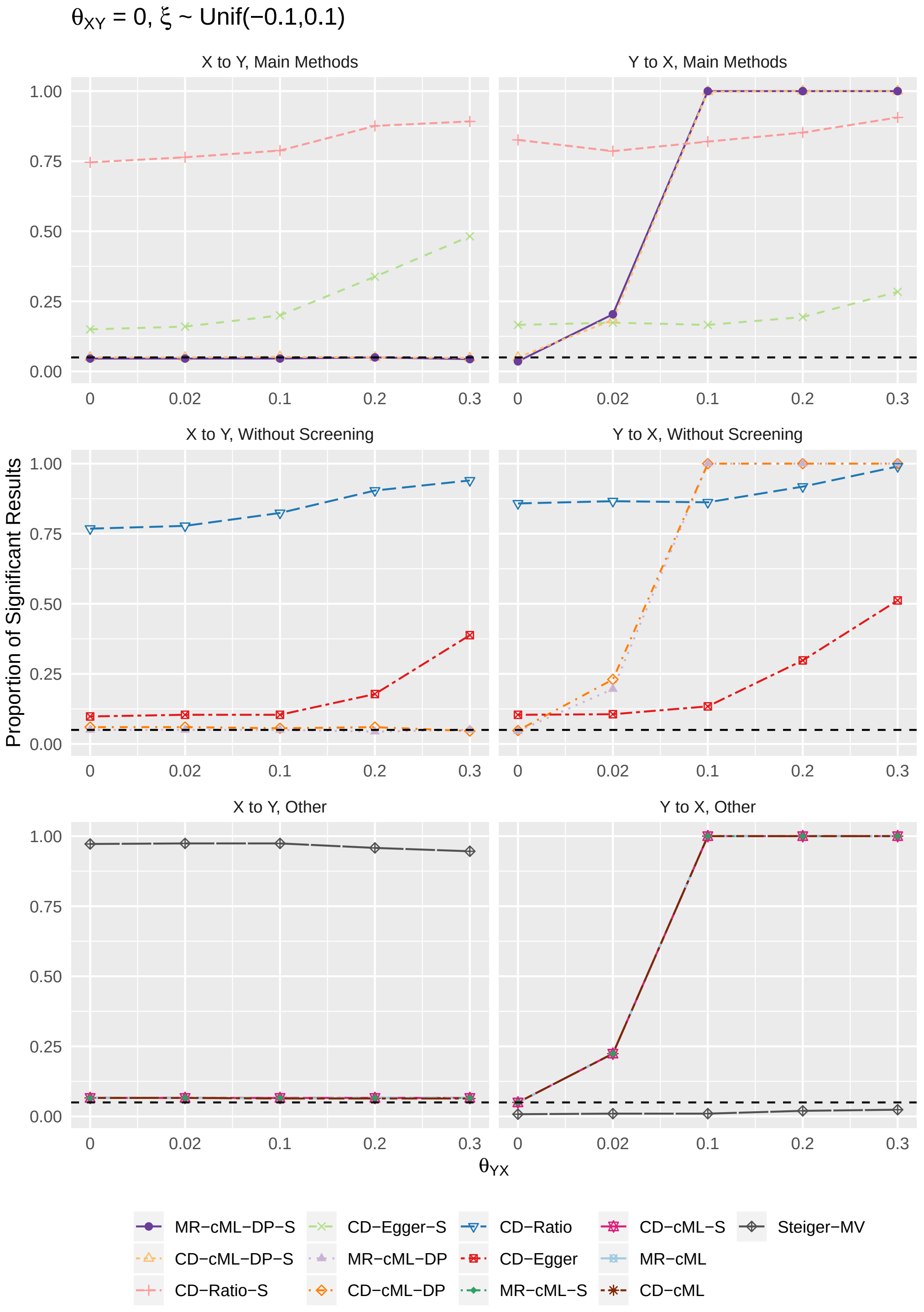

Figure S3: When $\theta_{XY} = 0, \xi \sim$ Unif(-0.2,0.2), proportions of significant simulation results obtained by each methods for direction $X \rightarrow Y$ (left column) and $Y \rightarrow X$ (right column). The first row shows results for four main methods: MR-cML-DP-S, CD-cML-DP-S, CD-Ratio-S, and CD-Egger-S; the second row shows results for four methods without screening: MR-cML-DP, CD-cML-DP, CD-Ratio, and CD-Egger; the third row shows results for other five methods.

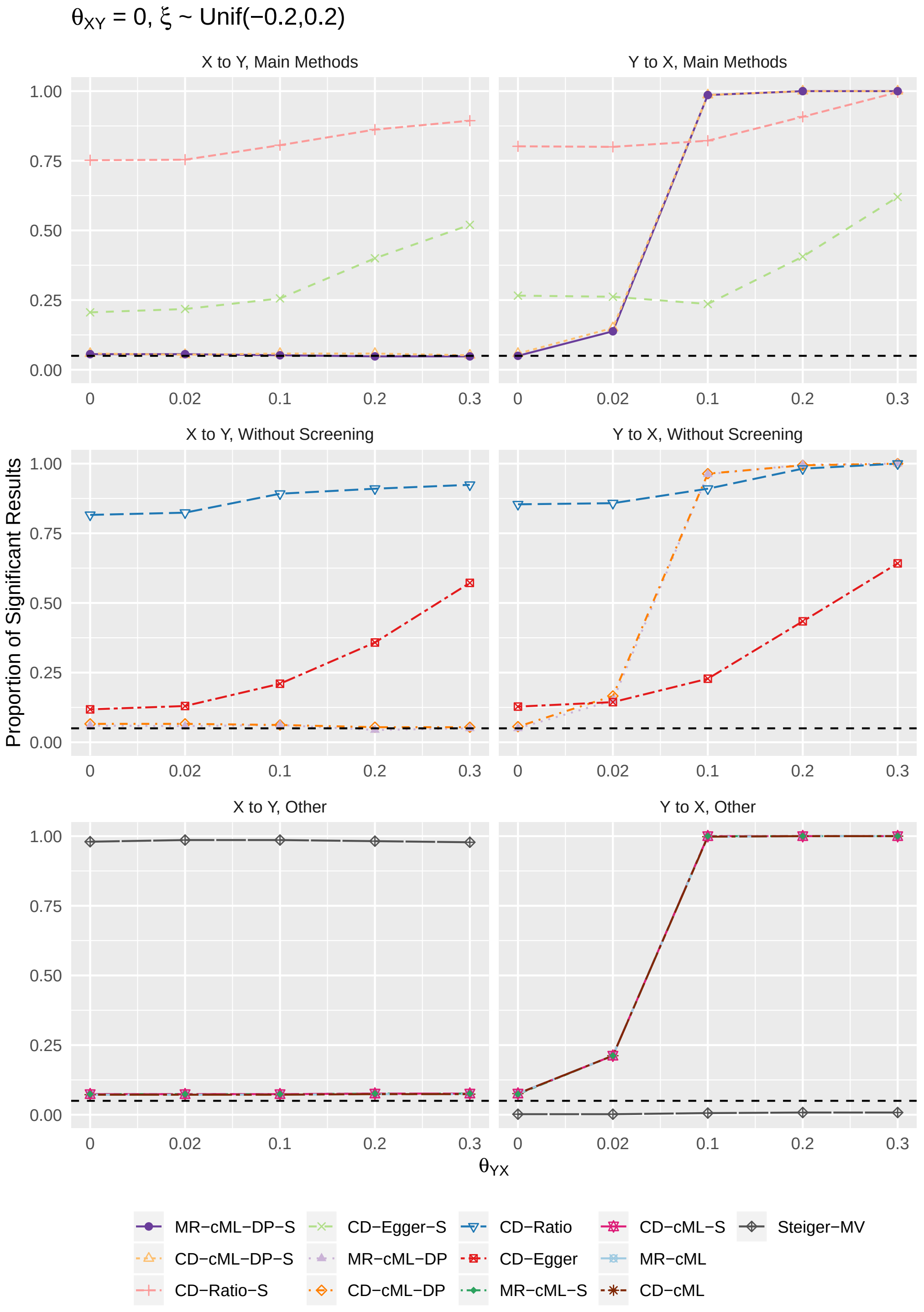

Figure S4: When $\theta_{XY} = 0.02, \xi = 0$, proportions of significant simulation results obtained by each methods for direction $X \to Y$ (left column) and $Y \to X$ (right column). The first row shows results for four main methods: MR-cML-DP-S, CD-cML-DP-S, CD-Ratio-S, and CD-Egger-S; the second row shows results for four methods without screening: MR-cML-DP, CD-cML-DP, CD-Ratio, and CD-Egger; the third row shows results for other five methods.

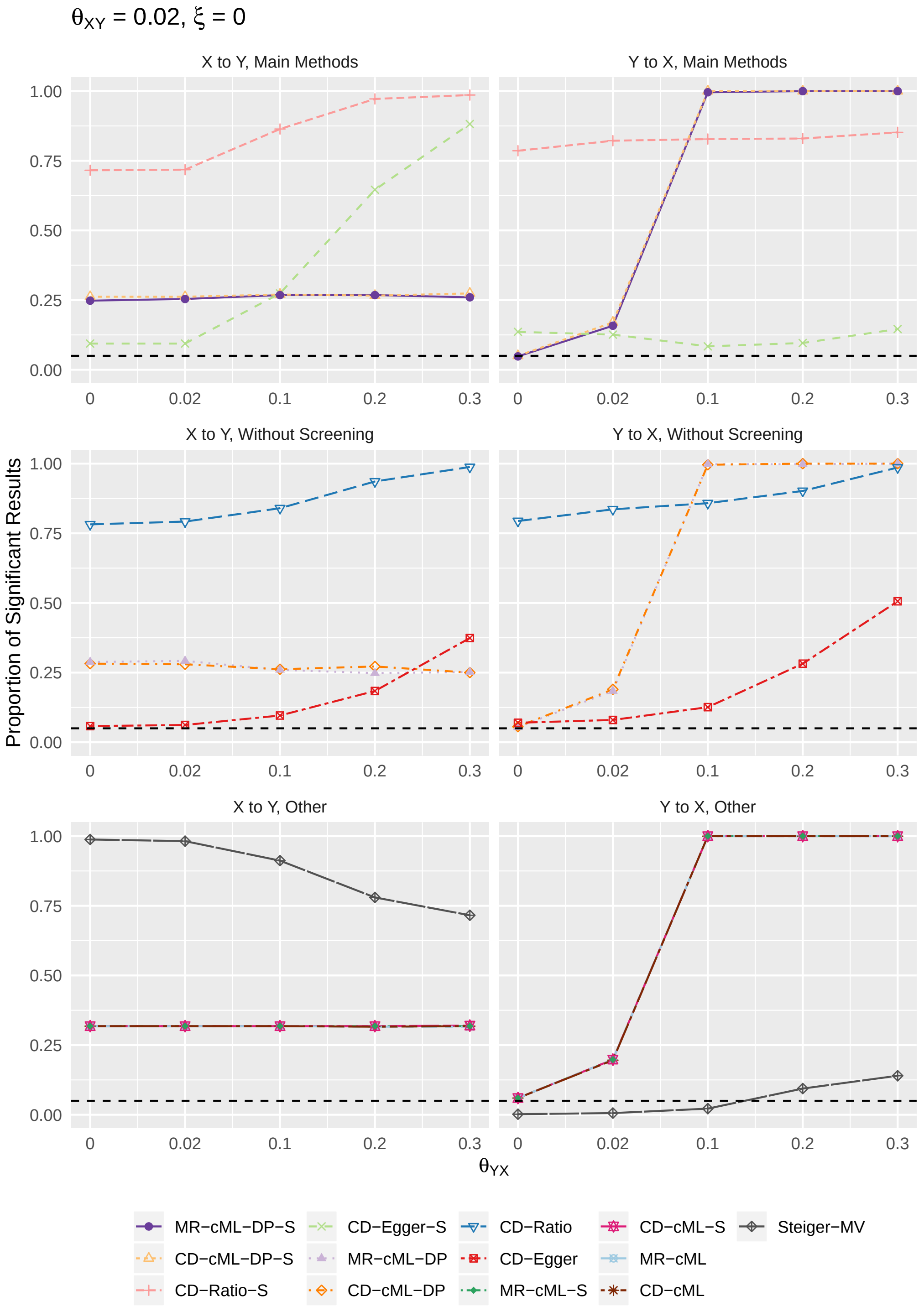

Figure S5: When $\theta_{XY} = 0.02, \xi \sim$ Unif(-0.1,0.1), proportions of significant simulation results obtained by each methods for direction $X \to Y$ (left column) and $Y \to X$ (right column). The first row shows results for four main methods: MR-cML-DP-S, CD-cML-DP-S, CD-Ratio-S, and CD-Egger-S; the second row shows results for four methods without screening: MR-cML-DP, CD-cML-DP, CD-Ratio, and CD-Egger; the third row shows results for other five methods.

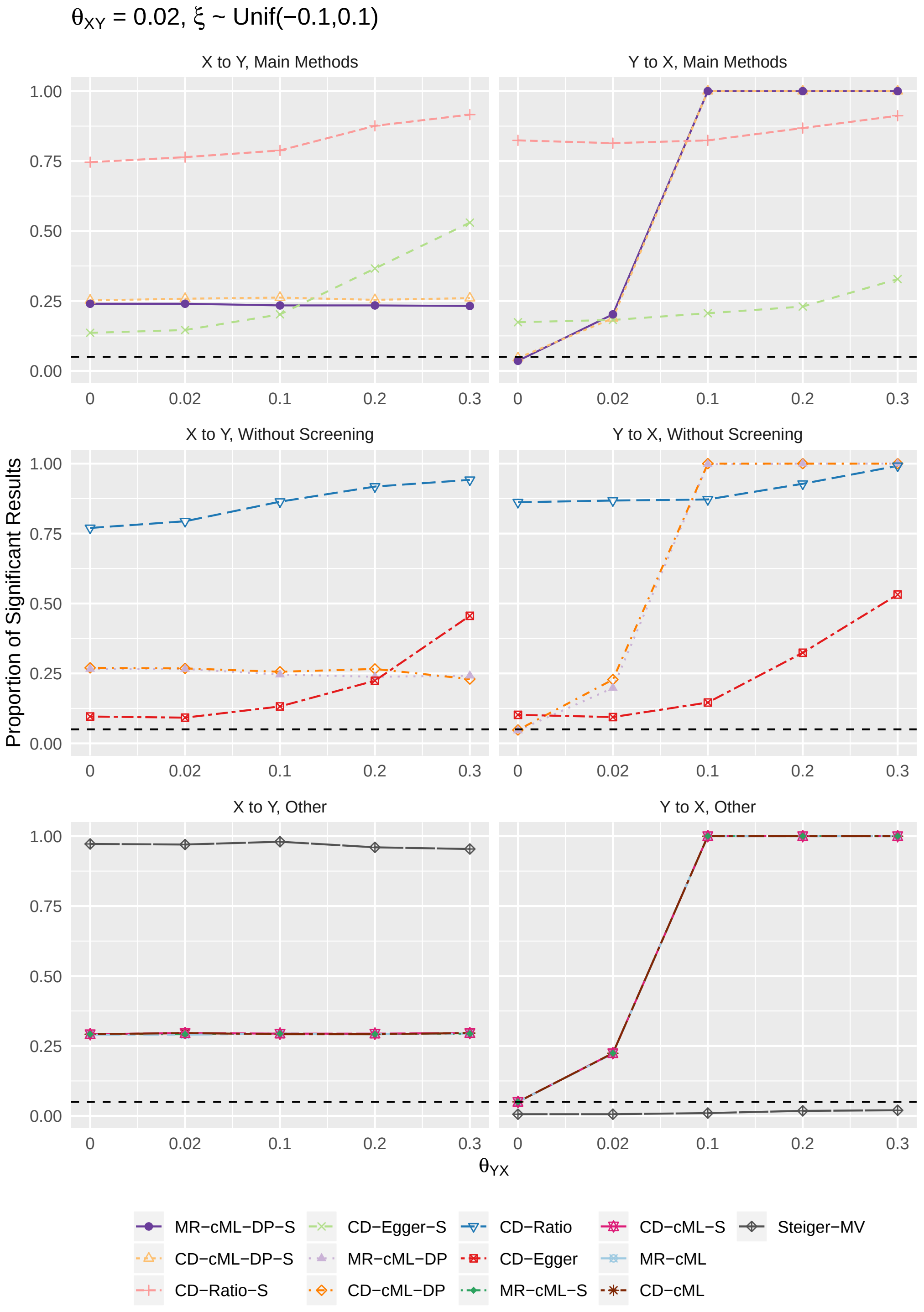

Figure S6: When $\theta_{XY} = 0.02, \xi \sim$ Unif(-0.2,0.2), proportions of significant simulation results obtained by each methods for direction $X \to Y$ (left column) and $Y \to X$ (right column). The first row shows results for four main methods: MR-cML-DP-S, CD-cML-DP-S, CD-Ratio-S, and CD-Egger-S; the second row shows results for four methods without screening: MR-cML-DP, CD-cML-DP, CD-Ratio, and CD-Egger; the third row shows results for other five methods.

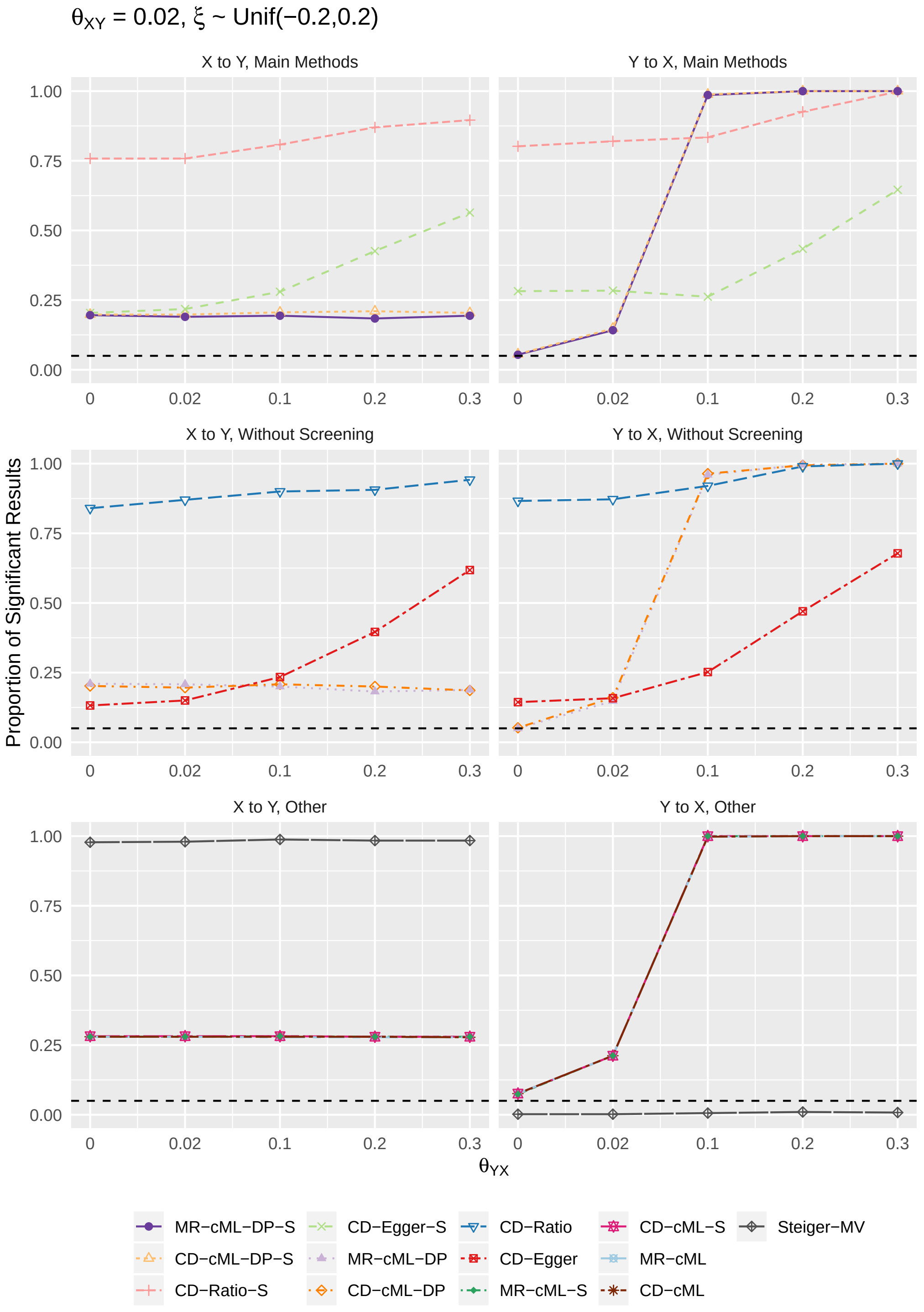

Figure S7: When $\theta_{XY} = 0.1, \xi = 0$, proportions of significant simulation results obtained by each methods for direction $X \to Y$ (left column) and $Y \to X$ (right column). The first row shows results for four main methods: MR-cML-DP-S, CD-cML-DP-S, CD-Ratio-S, and CD-Egger-S; the second row shows results for four methods without screening: MR-cML-DP, CD-cML-DP, CD-Ratio, and CD-Egger; the third row shows results for other five methods.

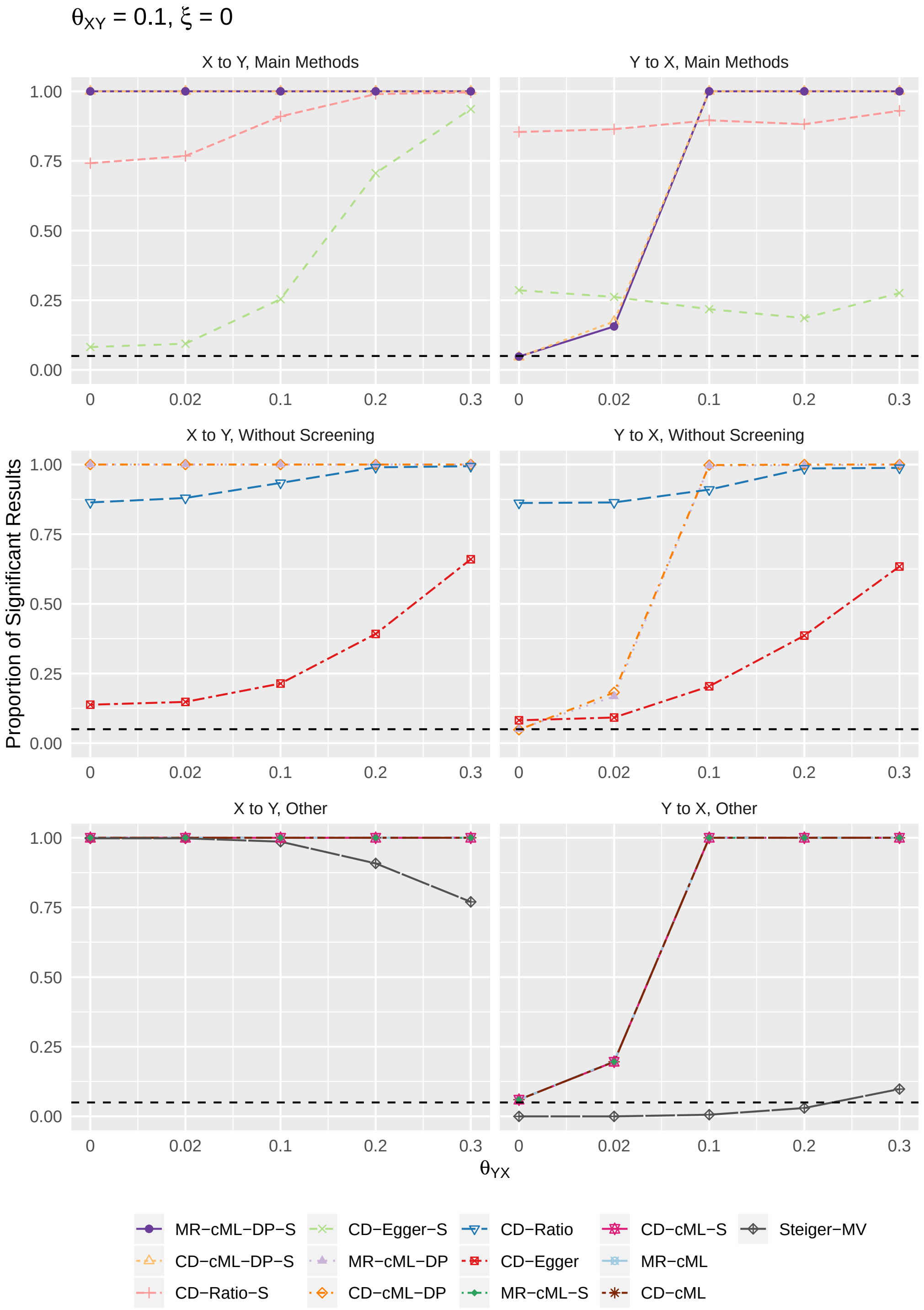

Figure S8: When $\theta_{XY} = 0.1, \xi \sim$ Unif(-0.1,0.1), proportions of significant simulation results obtained by each methods for direction $X \to Y$ (left column) and $Y \to X$ (right column). The first row shows results for four main methods: MR-cML-DP-S, CD-cML-DP-S, CD-Ratio-S, and CD-Egger-S; the second row shows results for four methods without screening: MR-cML-DP, CD-cML-DP, CD-Ratio, and CD-Egger; the third row shows results for other five methods.

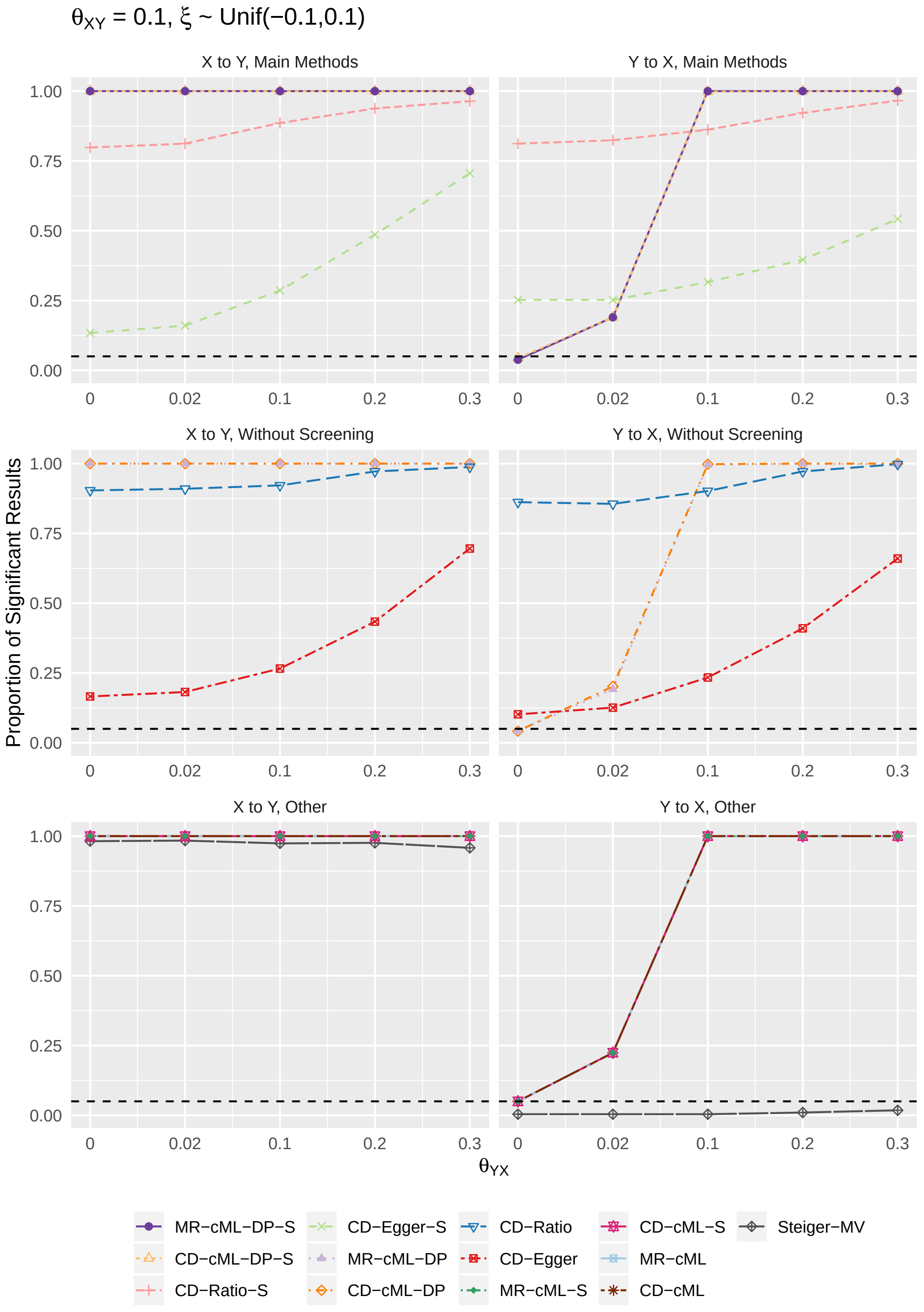

Figure S9: When $\theta_{XY} = 0.1, \xi \sim$ Unif(-0.2,0.2), proportions of significant simulation results obtained by each methods for direction $X \rightarrow Y$ (left column) and $Y \rightarrow X$ (right column). The first row shows results for four main methods: MR-cML-DP-S, CD-cML-DP-S, CD-Ratio-S, and CD-Egger-S; the second row shows results for four methods without screening: MR-cML-DP, CD-cML-DP, CD-Ratio, and CD-Egger; the third row shows results for other five methods.

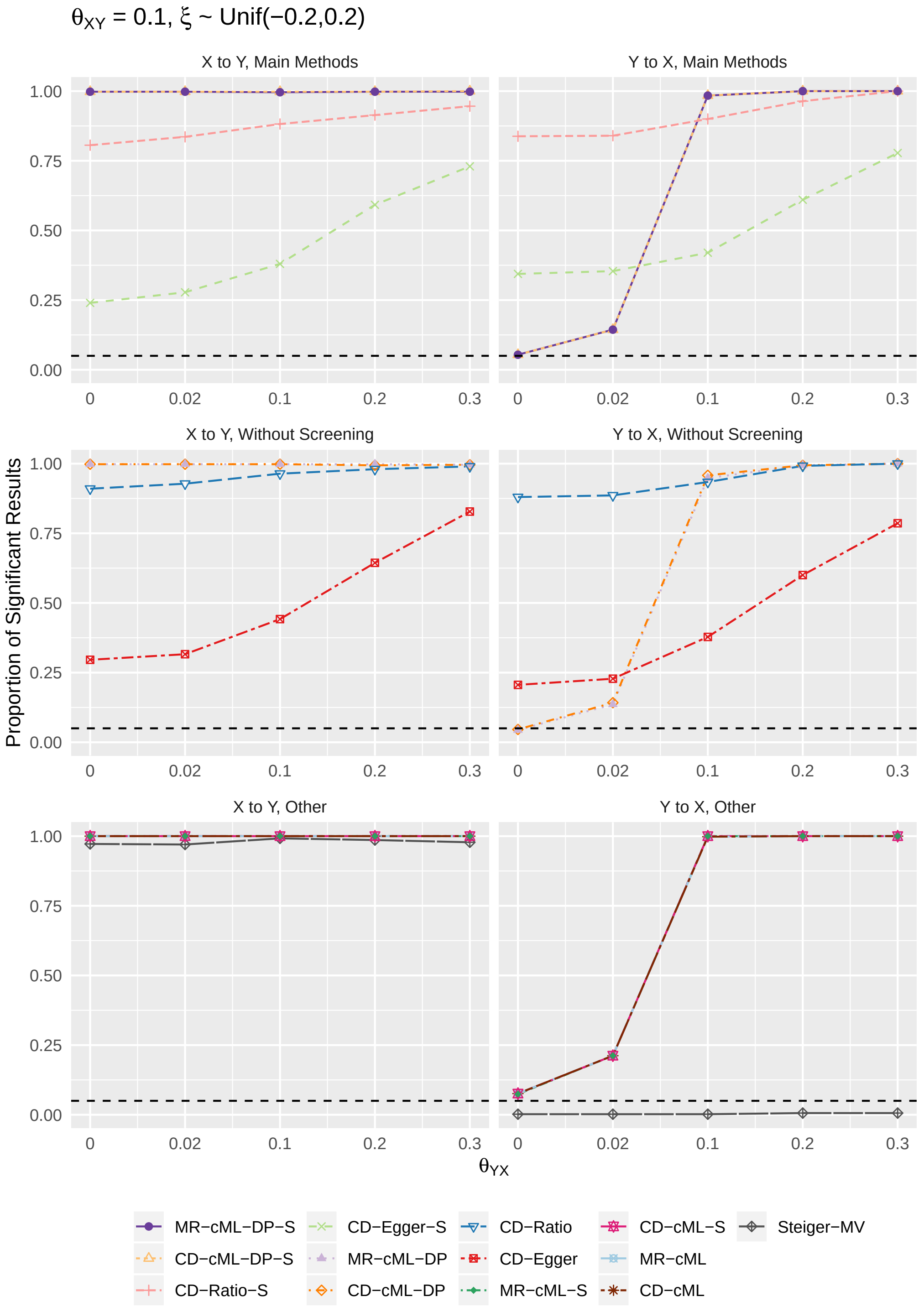

Figure S10: When $\theta_{XY} = 0.2, \xi = 0$, proportions of significant simulation results obtained by each methods for direction $X \rightarrow Y$ (left column) and $Y \rightarrow X$ (right column). The first row shows results for four main methods: MR-cML-DP-S, CD-cML-DP-S, CD-Ratio-S, and CD-Egger-S; the second row shows results for four methods without screening: MR-cML-DP, CD-cML-DP, CD-Ratio, and CD-Egger; the third row shows results for other five methods.

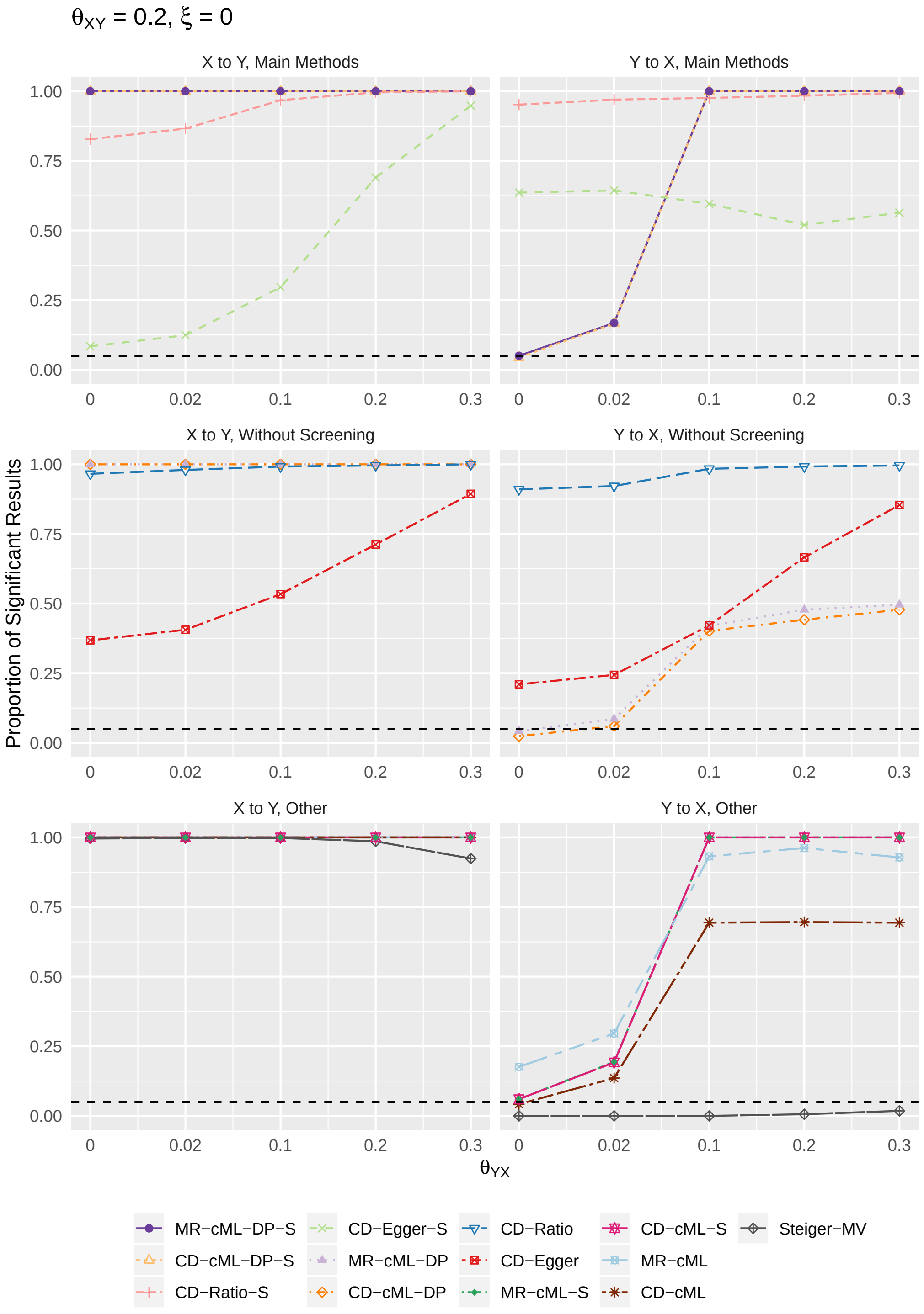

Figure S11: When $\theta_{XY} = 0.2, \xi \sim$ Unif(-0.1,0.1), proportions of significant simulation results obtained by each methods for direction $X \to Y$ (left column) and $Y \to X$ (right column). The first row shows results for four main methods: MR-cML-DP-S, CD-cML-DP-S, CD-Ratio-S, and CD-Egger-S; the second row shows results for four methods without screening: MR-cML-DP, CD-cML-DP, CD-Ratio, and CD-Egger; the third row shows results for other five methods.

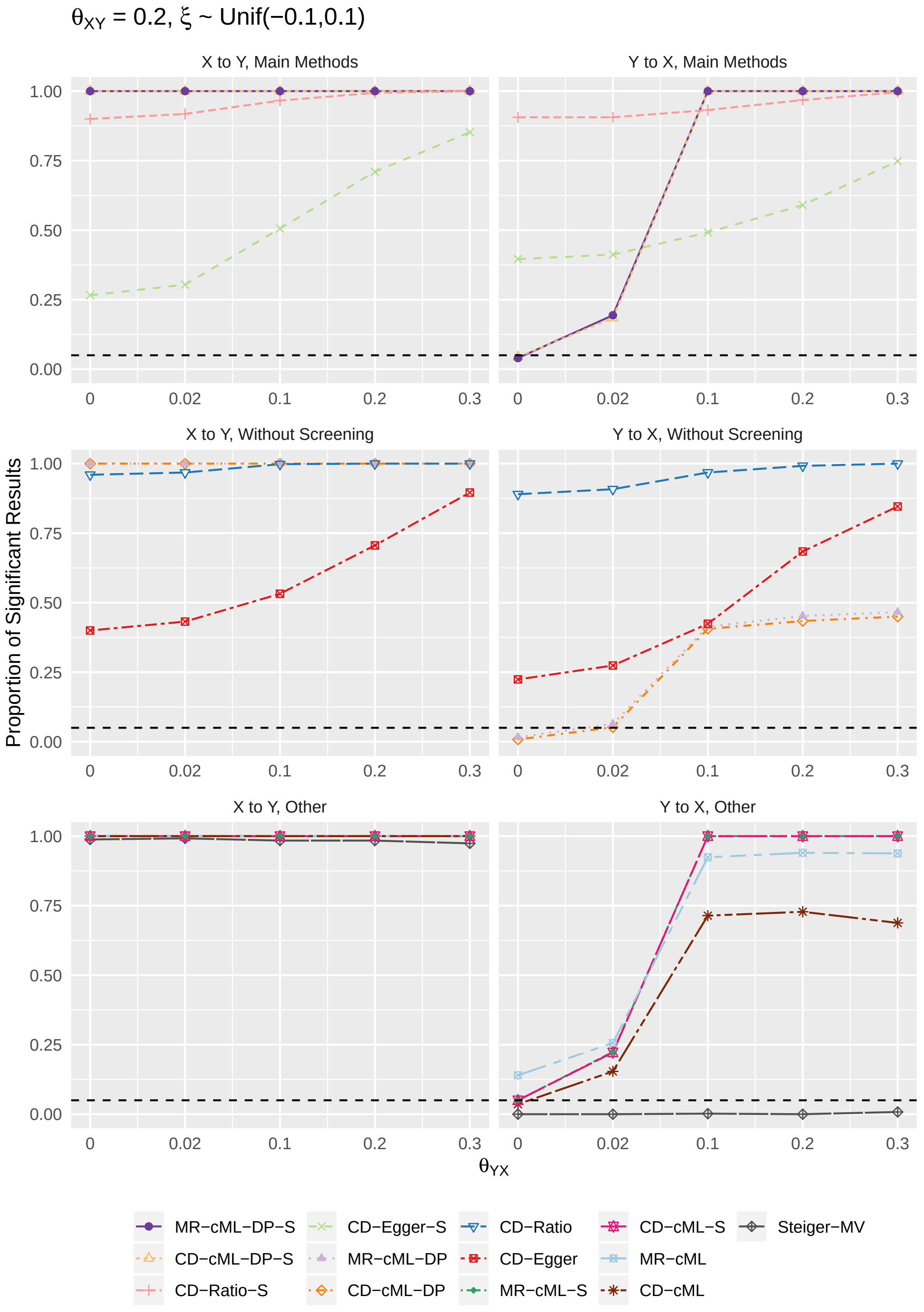

Figure S12: When $\theta_{XY} = 0.2, \xi \sim$ Unif(-0.2,0.2), proportions of significant simulation results obtained by each methods for direction $X \to Y$ (left column) and $Y \to X$ (right column). The first row shows results for four main methods: MR-cML-DP-S, CD-cML-DP-S, CD-Ratio-S, and CD-Egger-S; the second row shows results for four methods without screening: MR-cML-DP, CD-cML-DP, CD-Ratio, and CD-Egger; the third row shows results for other five methods.

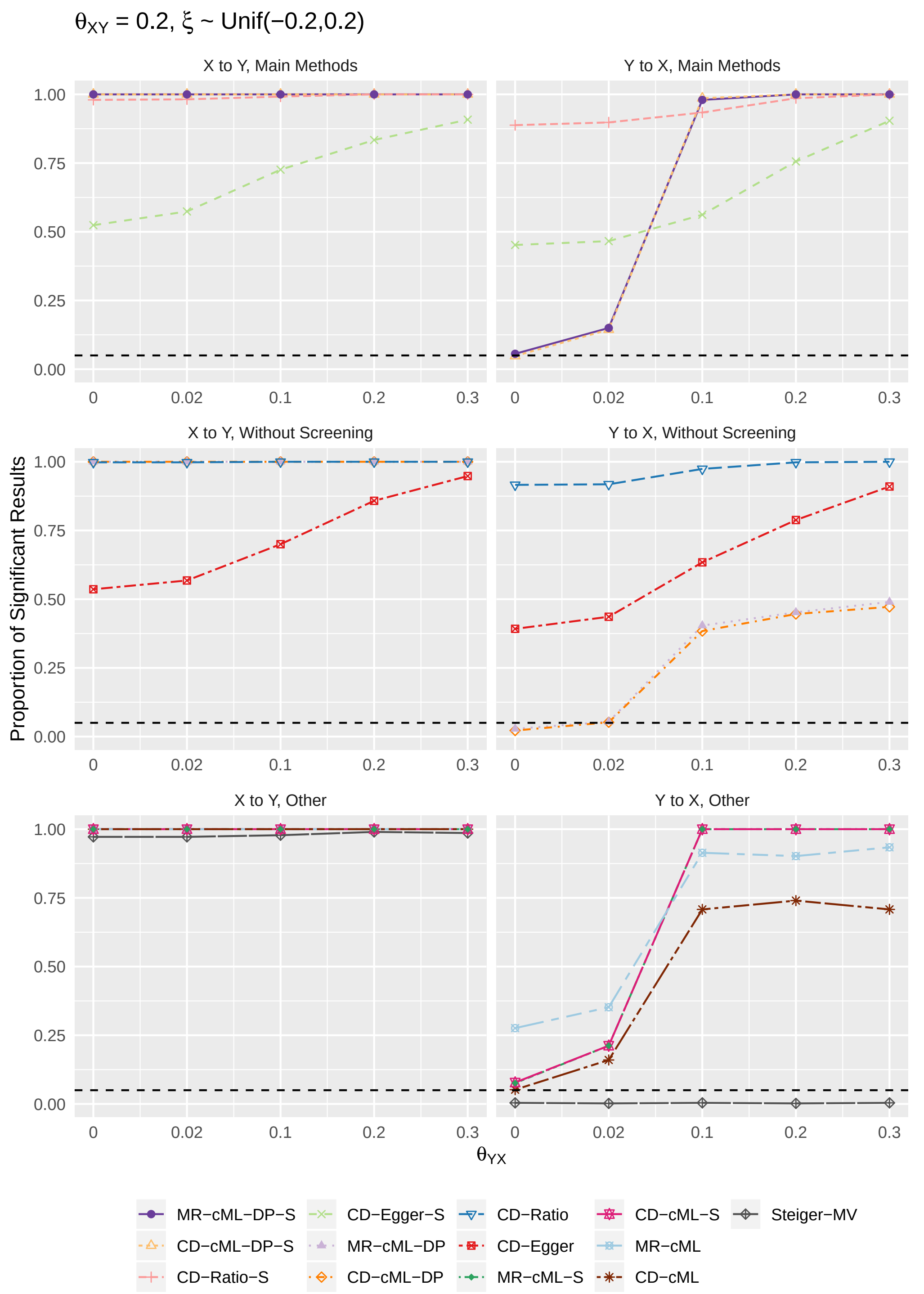

Figure S13: When $\theta_{XY} = 0.3, \xi = 0$, proportions of significant simulation results obtained by each methods for direction $X \to Y$ (left column) and $Y \to X$ (right column). The first row shows results for four main methods: MR-cML-DP-S, CD-cML-DP-S, CD-Ratio-S, and CD-Egger-S; the second row shows results for four methods without screening: MR-cML-DP, CD-cML-DP, CD-Ratio, and CD-Egger; the third row shows results for other five methods.

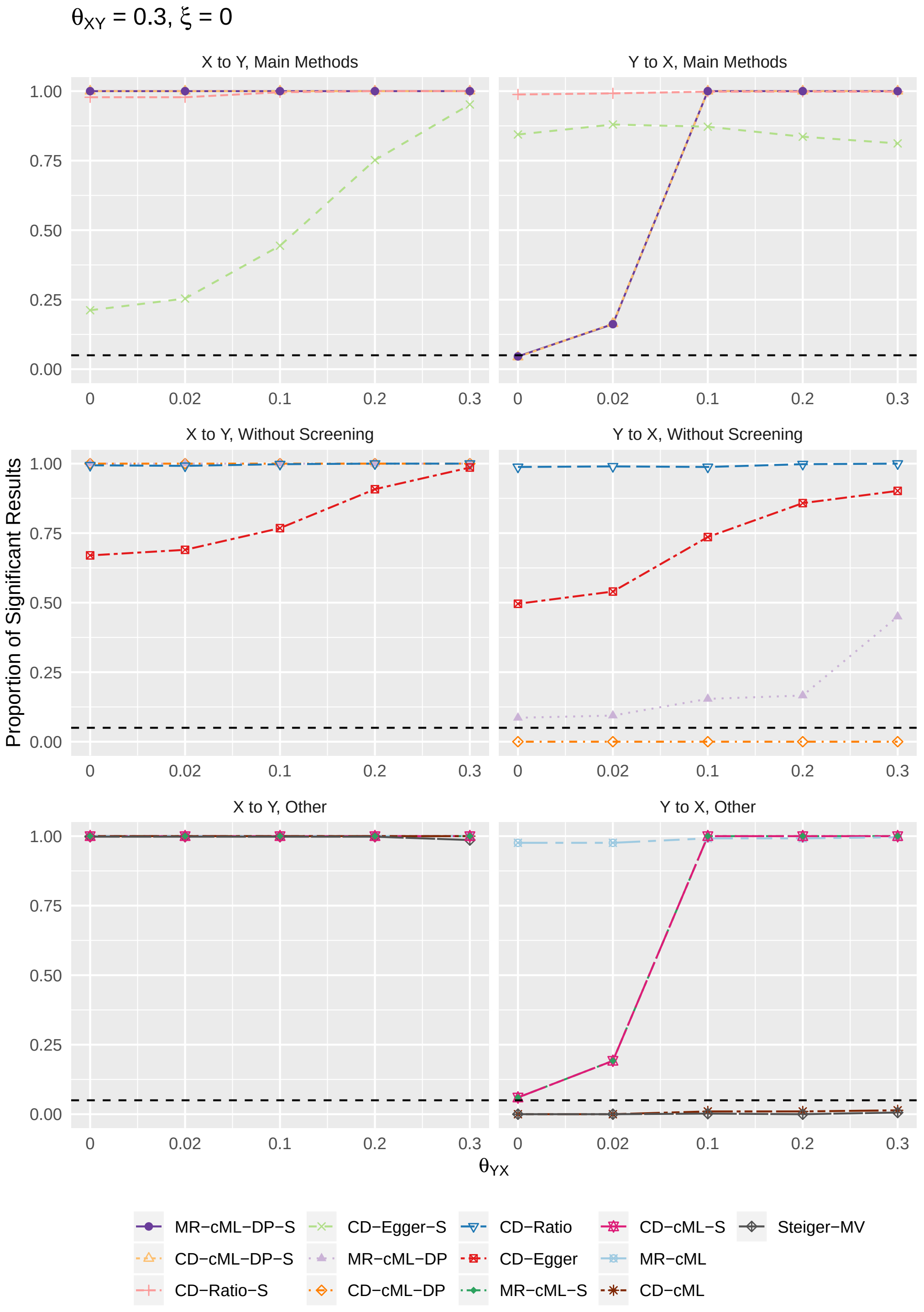

Figure S14: When $\theta_{XY} = 0.3, \xi \sim$ Unif(-0.1,0.1), proportions of significant simulation results obtained by each methods for direction $X \to Y$ (left column) and $Y \to X$ (right column). The first row shows results for four main methods: MR-cML-DP-S, CD-cML-DP-S, CD-Ratio-S, and CD-Egger-S; the second row shows results for four methods without screening: MR-cML-DP, CD-cML-DP, CD-Ratio, and CD-Egger; the third row shows results for other five methods.

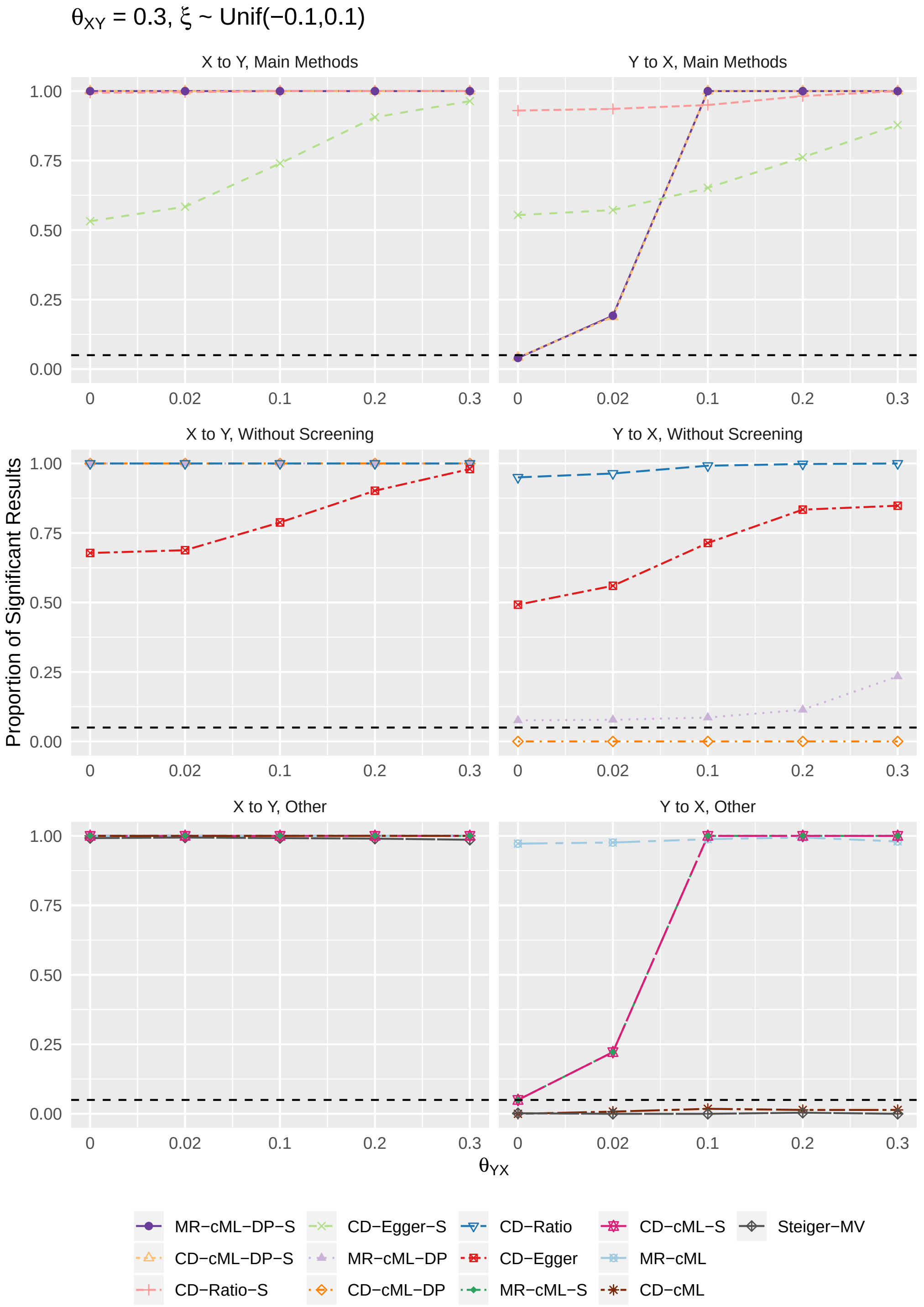

Figure S15: When $\theta_{XY} = 0.3, \xi \sim$ Unif(-0.2,0.2), proportions of significant simulation results obtained by each methods for direction $X \to Y$ (left column) and $Y \to X$ (right column). The first row shows results for four main methods: MR-cML-DP-S, CD-cML-DP-S, CD-Ratio-S, and CD-Egger-S; the second row shows results for four methods without screening: MR-cML-DP, CD-cML-DP, CD-Ratio, and CD-Egger; the third row shows results for other five methods.

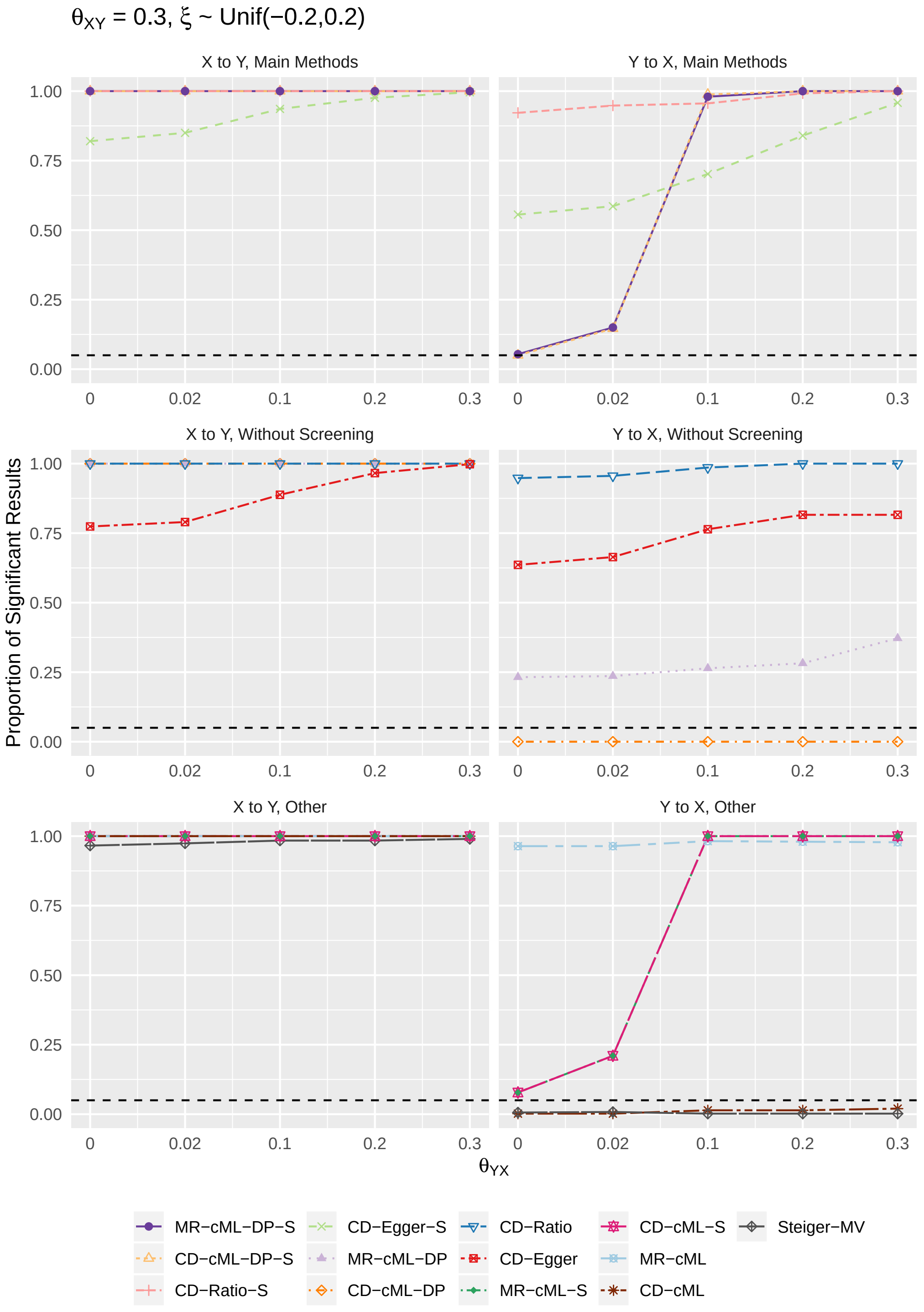

### S2 Full Real Data Results

#### S2.1 48 Exposure-Outcome Pairs

Table S1: Inferring causal effects between first 6 risk factors and CAD, in each cell we show the Bonferroni adjusted 1-0.05/48 $\approx$ 0.999 confidence interval of the estimate $\hat{\theta}$ for MR methods, and estimate $\hat{K}$ for CD methods; for Steiger's method, we show proportion of SNPs that give significant result. TRUE/FALSE in each cell indicates whether the result is significant or not, and cells give significant results are marked with red.

| Method \ Direction | TG to CAD | CAD to TG | LDL to CAD | CAD to LDL | HDL to CAD | CAD to HDL | Height to CAD | CAD to Height | BMI to CAD | CAD to BMI | BF to CAD | CAD to BF |
|---|---|---|---|---|---|---|---|---|---|---|---|---|
| MR-cML-DP-S | (0.134, 0.551), TRUE | (-0.046, 0.075), FALSE | (0.197, 0.657), TRUE | (-0.097, 0.027), FALSE | (-0.541, 0.018), FALSE | (-0.055, 0.052), FALSE | (-0.163, -0.042), TRUE | (-0.019, 0.066), FALSE | (0.149, 0.481), TRUE | (-0.081, 0.001), FALSE | (-0.631, 0.904), FALSE | (-0.101, 0.015), FALSE |
| MR-cML-S | (0.284, 0.456), TRUE | (-0.02, 0.045), FALSE | (0.293, 0.493), TRUE | (-0.074, 0.011), FALSE | (-0.341, -0.183), TRUE | (-0.037, 0.037), FALSE | (-0.148, -0.052), TRUE | (-0.006, 0.052), FALSE | (0.178, 0.419), TRUE | (-0.066, -0.011), TRUE | (-0.139, 0.557), FALSE | (-0.078, 0.001), FALSE |
| CD-cML-DP-S | (0.022, 0.118), TRUE | (-0.146, 0.256), FALSE | (0.056, 0.156), TRUE | (-0.373, 0.102), FALSE | (-0.11, 0.002), FALSE | (-0.182, 0.202), FALSE | (-0.047, -0.012), TRUE | (-0.059, 0.204), FALSE | (0.05, 0.157), TRUE | (-0.199, -0.004), TRUE | (-0.134, 0.213), FALSE | (-0.337, 0.052), FALSE |
| CD-cML-S | (0.056, 0.092), TRUE | (-0.083, 0.179), FALSE | (0.097, 0.133), TRUE | (-0.288, 0.045), FALSE | (-0.073, -0.039), TRUE | (-0.123, 0.144), FALSE | (-0.043, -0.015), TRUE | (-0.027, 0.169), FALSE | (0.059, 0.136), TRUE | (-0.171, -0.026), TRUE | (-0.031, 0.138), FALSE | (-0.261, 0.005), FALSE |
| CD-Ratio-S | (0.047, 0.077), TRUE | (-0.049, 0.204), FALSE | (0.08, 0.106), TRUE | (-0.008, 0.26), FALSE | (-0.052, -0.024), TRUE | (-0.151, 0.108), FALSE | (-0.037, -0.01), TRUE | (-0.063, 0.118), FALSE | (0.054, 0.129), TRUE | (-0.157, -0.018), TRUE | (-0.05, 0.089), FALSE | (-0.245, 0.015), FALSE |
| CD-Egger-S | (0.038, 0.104), TRUE | (-0.132, 0.326), FALSE | (0.063, 0.118), TRUE | (-0.133, 0.495), FALSE | (-0.09, -0.002), TRUE | (-0.269, 0.161), FALSE | (-0.047, -0.005), TRUE | (-0.194, 0.167), FALSE | (0.036, 0.161), TRUE | (-0.206, 0.04), FALSE | (-0.16, 0.136), FALSE | (-0.276, 0.074), FALSE |
| MR-cML-DP | (0.134, 0.551), TRUE | (-0.044, 0.078), FALSE | (0.19, 0.684), TRUE | (-0.095, 0.027), FALSE | (-0.541, 0.018), FALSE | (-0.052, 0.051), FALSE | (-0.163, -0.042), TRUE | (-0.021, 0.071), FALSE | (0.149, 0.481), TRUE | (-0.081, 0.001), FALSE | (-0.631, 0.904), FALSE | (-0.101, 0.015), FALSE |
| MR-cML | (0.284, 0.456), TRUE | (-0.02, 0.045), FALSE | (0.235, 0.574), TRUE | (-0.074, 0.011), FALSE | (-0.341, -0.183), TRUE | (-0.037, 0.037), FALSE | (-0.148, -0.052), TRUE | (-0.006, 0.052), FALSE | (0.178, 0.419), TRUE | (-0.066, -0.011), TRUE | (-0.139, 0.557), FALSE | (-0.078, 0.001), FALSE |
| CD-cML-DP | (0.022, 0.118), TRUE | (-0.146, 0.269), FALSE | (0.056, 0.159), TRUE | (-0.349, 0.092), FALSE | (-0.11, 0.002), FALSE | (-0.166, 0.19), FALSE | (-0.047, -0.012), TRUE | (-0.067, 0.214), FALSE | (0.05, 0.157), TRUE | (-0.199, -0.004), TRUE | (-0.134, 0.213), FALSE | (-0.337, 0.052), FALSE |
| CD-cML | (0.056, 0.092), TRUE | (-0.083, 0.179), FALSE | (0.098, 0.134), TRUE | (-0.288, 0.045), FALSE | (-0.073, -0.039), TRUE | (-0.123, 0.144), FALSE | (-0.043, -0.015), TRUE | (-0.027, 0.169), FALSE | (0.059, 0.136), TRUE | (-0.171, -0.026), TRUE | (-0.031, 0.138), FALSE | (-0.261, 0.005), FALSE |
| CD-Ratio | (0.047, 0.077), TRUE | (-0.009, 0.242), FALSE | (0.082, 0.107), TRUE | (0.057, 0.32), TRUE | (-0.052, -0.024), TRUE | (-0.296, -0.047), TRUE | (-0.037, -0.01), TRUE | (-0.074, 0.107), FALSE | (0.054, 0.129), TRUE | (-0.157, -0.018), TRUE | (-0.05, 0.089), FALSE | (-0.245, 0.015), FALSE |
| CD-Egger | (0.038, 0.104), TRUE | (-0.128, 0.833), FALSE | (0.056, 0.15), TRUE | (-0.042, 2.15), FALSE | (-0.09, -0.002), TRUE | (-1.009, 0.112), FALSE | (-0.047, -0.005), TRUE | (-0.195, 0.19), FALSE | (0.036, 0.161), TRUE | (-0.206, 0.04), FALSE | (-0.16, 0.136), FALSE | (-0.276, 0.074), FALSE |
| Steiger | 0.505, TRUE | 0.184, FALSE | 0.608, TRUE | 0.128, FALSE | 0.617, TRUE | 0.141, FALSE | 0.914, TRUE | 0.073, FALSE | 0.492, TRUE | 0.407, FALSE | 0.111, FALSE | 0.25, FALSE |

Table S2: Inferring causal effects between second 6 risk factors and CAD, in each cell we show the Bonferroni adjusted 1-0.05/48 $\approx$ 0.999 confidence interval of the estimate $\hat{\theta}$ for MR methods, and estimate $\hat{K}$ for CD methods; for Steiger's method, we show proportion of SNPs that give significant result. TRUE/FALSE in each cell indicates whether the result is significant or not, and cells give significant results are marked with red.

| Direction / Method | BW to CAD | CAD to BW | DBP to CAD | CAD to DBP | SBP to CAD | CAD to SBP | FG to CAD | CAD to FG | Smoke to CAD | CAD to Smoke | Alcohol to CAD | CAD to Alcohol |
|---|---|---|---|---|---|---|---|---|---|---|---|---|
| MR-cML-DP-S | (-0.356, 0.138), FALSE | (-0.061, 0.024), FALSE | (0.056, 0.086), TRUE | (-0.921, 1.016), FALSE | (0.037, 0.053), TRUE | (-1.339, 4.208), FALSE | (0.138, 0.546), TRUE | (-0.026, 0.048), FALSE | (-0.151, 0.363), FALSE | (-0.069, 0.034), FALSE | (-0.185, 0.753), FALSE | (-0.013, 0.022), FALSE |
| MR-cML-S | (-0.255, 0.019), FALSE | (-0.048, 0.007), FALSE | (0.058, 0.074), TRUE | (-0.237, 0.137), FALSE | (0.04, 0.049), TRUE | (-0.216, 4.24), FALSE | (0.151, 0.536), TRUE | (-0.018, 0.038), FALSE | (-0.081, 0.3), FALSE | (-0.055, 0.024), FALSE | (0.032, 0.543), TRUE | (-0.01, 0.018), FALSE |
| CD-cML-DP-S | (-0.098, 0.039), FALSE | (-0.221, 0.089), FALSE | (0.163, 0.25), TRUE | (-0.309, 0.342), FALSE | (0.185, 0.262), TRUE | (-0.27, 0.83), FALSE | (0.021, 0.085), TRUE | (-0.174, 0.293), FALSE | (-0.081, 0.196), FALSE | (-0.126, 0.064), FALSE | (-0.053, 0.211), FALSE | (-0.049, 0.082), FALSE |
| CD-cML-S | (-0.07, 0.005), FALSE | (-0.176, 0.028), FALSE | (0.169, 0.216), TRUE | (-0.079, 0.052), FALSE | (0.2, 0.246), TRUE | (-0.239, 0.939), FALSE | (0.023, 0.083), TRUE | (-0.122, 0.233), FALSE | (-0.043, 0.163), FALSE | (-0.1, 0.046), FALSE | (0.009, 0.152), TRUE | (-0.035, 0.067), FALSE |
| CD-Ratio-S | (-0.063, 0.007), FALSE | (-0.175, 0.019), FALSE | (0.152, 0.194), TRUE | (-0.019, 0.085), FALSE | (0.172, 0.215), TRUE | (0.072, 0.173), TRUE | (0.014, 0.073), TRUE | (-0.125, 0.227), FALSE | (-0.024, 0.076), FALSE | (-0.09, 0.05), FALSE | (-0.014, 0.119), FALSE | (-0.027, 0.07), FALSE |
| CD-Egger-S | (-0.114, 0.034), FALSE | (-0.237, 0.048), FALSE | (0.144, 0.224), TRUE | (-0.171, 0.238), FALSE | (0.17, 0.249), TRUE | (-0.032, 0.317), FALSE | (-0.006, 0.105), FALSE | (-0.175, 0.278), FALSE | (-0.039, 0.11), FALSE | (-0.129, 0.113), FALSE | (-0.09, 0.179), FALSE | (-0.048, 0.09), FALSE |
| MR-cML-DP | (-0.356, 0.138), FALSE | (-0.063, 0.029), FALSE | (0.055, 0.087), TRUE | (-1.906, 2.258), FALSE | (0.037, 0.052), TRUE | (-2.054, 4.805), FALSE | (0.138, 0.546), TRUE | (-0.026, 0.048), FALSE | (-0.151, 0.363), FALSE | (-0.069, 0.034), FALSE | (-0.185, 0.753), FALSE | (-0.013, 0.022), FALSE |
| MR-cML | (-0.255, 0.019), FALSE | (-0.048, 0.007), FALSE | (0.058, 0.074), TRUE | (-0.237, 0.137), FALSE | (0.04, 0.049), TRUE | (1.945, 3.012), TRUE | (0.151, 0.536), TRUE | (-0.018, 0.038), FALSE | (-0.081, 0.3), FALSE | (-0.055, 0.024), FALSE | (0.032, 0.543), TRUE | (-0.01, 0.018), FALSE |
| CD-cML-DP | (-0.098, 0.039), FALSE | (-0.224, 0.104), FALSE | (0.161, 0.252), TRUE | (-0.598, 0.691), FALSE | (0.186, 0.261), TRUE | (-0.422, 0.961), FALSE | (0.021, 0.085), TRUE | (-0.174, 0.293), FALSE | (-0.081, 0.196), FALSE | (-0.126, 0.064), FALSE | (-0.053, 0.211), FALSE | (-0.049, 0.082), FALSE |
| CD-cML | (-0.07, 0.005), FALSE | (-0.176, 0.028), FALSE | (0.169, 0.216), TRUE | (-0.079, 0.052), FALSE | (0.2, 0.246), TRUE | (0.357, 0.612), TRUE | (0.023, 0.083), TRUE | (-0.122, 0.233), FALSE | (-0.043, 0.163), FALSE | (-0.1, 0.046), FALSE | (0.009, 0.152), TRUE | (-0.035, 0.067), FALSE |
| CD-Ratio | (-0.063, 0.007), FALSE | (-0.19, 0.003), FALSE | (0.153, 0.195), TRUE | (0.033, 0.134), TRUE | (0.174, 0.217), TRUE | (0.123, 0.222), TRUE | (0.014, 0.073), TRUE | (-0.125, 0.227), FALSE | (-0.024, 0.076), FALSE | (-0.09, 0.05), FALSE | (-0.014, 0.119), FALSE | (-0.027, 0.07), FALSE |
| CD-Egger | (-0.114, 0.034), FALSE | (-0.271, 0.04), FALSE | (0.144, 0.229), TRUE | (-0.164, 0.672), FALSE | (0.166, 0.255), TRUE | (0.055, 0.639), TRUE | (-0.006, 0.105), FALSE | (-0.175, 0.278), FALSE | (-0.039, 0.11), FALSE | (-0.129, 0.113), FALSE | (-0.09, 0.179), FALSE | (-0.048, 0.09), FALSE |
| Steiger | 0.407, TRUE | 0.296, FALSE | 0.833, TRUE | 0.055, FALSE | 0.809, TRUE | 0.071, FALSE | 0.167, FALSE | 0.097, FALSE | 0.209, FALSE | 0.64, TRUE | 0.27, FALSE | 0.69, TRUE |

Table S3: Inferring causal effects between first 6 risk factors and Stroke, in each cell we show the Bonferroni adjusted 1-0.05/48 $\approx$ 0.999 confidence interval of the estimate $\hat{\theta}$ for MR methods, and estimate $\hat{K}$ for CD methods; for Steiger's method, we show proportion of SNPs that give significant result. TRUE/FALSE in each cell indicates whether the result is significant or not, and cells give significant results are marked with red.

| Direction<br>Method | TG to Stroke | Stroke to TG | LDL to Stroke | Stroke to LDL | HDL to Stroke | Stroke to HDL | Height to Stroke | Stroke to Height | BMI to Stroke | Stroke to BMI | BF to Stroke | Stroke to BF |
|---|---|---|---|---|---|---|---|---|---|---|---|---|
| MR-cML-DP-S | (-0.103, 0.109), FALSE | (-0.023, 0.173), FALSE | (0.014, 0.194), TRUE | (-0.179, 0.131), FALSE | (-0.149, 0.024), FALSE | (-0.101, 0.105), FALSE | (-0.087, 0.058), FALSE | (-0.223, 0.127), FALSE | (-0.005, 0.29), FALSE | (-0.106, 0.103), FALSE | (-0.198, 0.432), FALSE | (-0.093, 0.098), FALSE |
| MR-cML-S | (-0.083, 0.083), FALSE | (-0.004, 0.151), FALSE | (0.048, 0.175), TRUE | (-0.135, 0.082), FALSE | (-0.134, 0.015), FALSE | (-0.082, 0.09), FALSE | (-0.065, 0.042), FALSE | (-0.155, 0.029), FALSE | (0.018, 0.285), TRUE | (-0.081, 0.081), FALSE | (-0.172, 0.383), FALSE | (-0.09, 0.087), FALSE |
| CD-cML-DP-S | (-0.021, 0.022), FALSE | (-0.127, 0.647), FALSE | (0.004, 0.044), TRUE | (-0.569, 0.403), FALSE | (-0.031, 0.004), FALSE | (-0.333, 0.423), FALSE | (-0.024, 0.017), FALSE | (-0.825, 0.462), FALSE | (-0.004, 0.091), FALSE | (-0.237, 0.252), FALSE | (-0.05, 0.11), FALSE | (-0.317, 0.348), FALSE |
| CD-cML-S | (-0.017, 0.016), FALSE | (-0.064, 0.582), FALSE | (0.011, 0.039), TRUE | (-0.435, 0.271), FALSE | (-0.028, 0.003), FALSE | (-0.274, 0.38), FALSE | (-0.018, 0.012), FALSE | (-0.577, 0.114), FALSE | (0.004, 0.089), TRUE | (-0.194, 0.209), FALSE | (-0.043, 0.098), FALSE | (-0.309, 0.314), FALSE |
| CD-Ratio-S | (-0.016, 0.017), FALSE | (-0.045, 0.544), FALSE | (0.009, 0.036), TRUE | (-0.342, 0.271), FALSE | (-0.029, 0.001), FALSE | (-0.276, 0.373), FALSE | (-0.023, 0.006), FALSE | (-0.37, 0.226), FALSE | (0, 0.082), TRUE | (-0.232, 0.142), FALSE | (-0.044, 0.097), FALSE | (-0.307, 0.311), FALSE |
| CD-Egger-S | (-0.021, 0.023), FALSE | (-0.101, 0.655), FALSE | (0.001, 0.045), TRUE | (-0.45, 0.447), FALSE | (-0.038, 0.005), FALSE | (-0.335, 0.452), FALSE | (-0.029, 0.011), FALSE | (-0.731, 0.446), FALSE | (-0.018, 0.1), FALSE | (-0.37, 0.221), FALSE | (-0.046, 0.097), FALSE | (-0.299, 0.313), FALSE |
| MR-cML-DP | (-0.103, 0.109), FALSE | (-0.023, 0.173), FALSE | (0.014, 0.194), TRUE | (-0.179, 0.128), FALSE | (-0.149, 0.024), FALSE | (-0.11, 0.128), FALSE | (-0.087, 0.058), FALSE | (-0.203, 0.111), FALSE | (-0.005, 0.29), FALSE | (-0.106, 0.103), FALSE | (-0.198, 0.432), FALSE | (-0.093, 0.098), FALSE |
| MR-cML | (-0.083, 0.083), FALSE | (-0.004, 0.151), FALSE | (0.048, 0.175), TRUE | (-0.135, 0.082), FALSE | (-0.134, 0.015), FALSE | (-0.082, 0.09), FALSE | (-0.065, 0.042), FALSE | (-0.155, 0.029), FALSE | (0.018, 0.285), TRUE | (-0.081, 0.081), FALSE | (-0.172, 0.383), FALSE | (-0.09, 0.087), FALSE |
| CD-cML-DP | (-0.021, 0.022), FALSE | (-0.127, 0.647), FALSE | (0.004, 0.044), TRUE | (-0.549, 0.382), FALSE | (-0.031, 0.004), FALSE | (-0.367, 0.505), FALSE | (-0.024, 0.017), FALSE | (-0.753, 0.417), FALSE | (-0.004, 0.091), FALSE | (-0.237, 0.252), FALSE | (-0.05, 0.11), FALSE | (-0.317, 0.348), FALSE |
| CD-cML | (-0.017, 0.016), FALSE | (-0.064, 0.582), FALSE | (0.011, 0.039), TRUE | (-0.435, 0.271), FALSE | (-0.028, 0.003), FALSE | (-0.274, 0.38), FALSE | (-0.018, 0.012), FALSE | (-0.577, 0.114), FALSE | (0.004, 0.089), TRUE | (-0.194, 0.209), FALSE | (-0.043, 0.098), FALSE | (-0.309, 0.314), FALSE |
| CD-Ratio | (-0.016, 0.017), FALSE | (-0.045, 0.544), FALSE | (0.009, 0.036), TRUE | (-0.424, 0.177), FALSE | (-0.029, 0.001), FALSE | (-0.371, 0.265), FALSE | (-0.023, 0.006), FALSE | (-0.421, 0.172), FALSE | (0, 0.082), TRUE | (-0.232, 0.142), FALSE | (-0.044, 0.097), FALSE | (-0.307, 0.311), FALSE |
| CD-Egger | (-0.021, 0.023), FALSE | (-0.101, 0.655), FALSE | (0.001, 0.045), TRUE | (-0.865, 0.413), FALSE | (-0.038, 0.005), FALSE | (-0.9, 0.512), FALSE | (-0.029, 0.011), FALSE | (-2.113, 0.848), FALSE | (-0.018, 0.1), FALSE | (-0.37, 0.221), FALSE | (-0.046, 0.097), FALSE | (-0.299, 0.313), FALSE |
| Steiger | 0.786, TRUE | 0.086, FALSE | 0.846, TRUE | 0.066, FALSE | 0.879, TRUE | 0.061, FALSE | 0.984, TRUE | 0.011, FALSE | 0.829, TRUE | 0.132, FALSE | 0.4, TRUE | 0.32, FALSE |

Table S4: Inferring causal effects between second 6 risk factors and Stroke, in each cell we show the Bonferroni adjusted 1-0.05/48 $\approx$ 0.999 confidence interval of the estimate $\hat{\theta}$ for MR methods, and estimate $\hat{K}$ for CD methods; for Steiger's method, we show proportion of SNPs that give significant result. TRUE/FALSE in each cell indicates whether the result is significant or not, and cells give significant results are marked with red.

| Direction / Method | BW to Stroke | Stroke to BW | DBP to Stroke | Stroke to DBP | SBP to Stroke | Stroke to SBP | FG to Stroke | Stroke to FG | Smoke to Stroke | Stroke to Smoke | Alcohol to Stroke | Stroke to Alcohol |
|---|---|---|---|---|---|---|---|---|---|---|---|---|
| MR-cML-DP-S | (-0.272, 0.235), FALSE | (-0.156, 0.081), FALSE | (0.043, 0.068), TRUE | (-1.276, -0.055), TRUE | (0.03, 0.044), TRUE | (-1.448, 5.095), FALSE | (-0.253, 0.634), FALSE | (-0.045, 0.109), FALSE | (-0.093, 0.229), FALSE | (-0.101, 0.142), FALSE | (-0.047, 0.498), FALSE | (-0.036, 0.031), FALSE |
| MR-cML-S | (-0.198, 0.136), FALSE | (-0.099, 0.046), FALSE | (0.046, 0.064), TRUE | (-1.205, -0.135), TRUE | (0.031, 0.041), TRUE | (0, 4.104), FALSE | (-0.069, 0.408), FALSE | (-0.03, 0.101), FALSE | (-0.038, 0.19), FALSE | (-0.081, 0.1), FALSE | (-0.015, 0.46), FALSE | (-0.032, 0.028), FALSE |
| CD-cML-DP-S | (-0.07, 0.058), FALSE | (-0.568, 0.303), FALSE | (0.118, 0.185), TRUE | (-0.476, -0.02), TRUE | (0.142, 0.21), TRUE | (-0.362, 1.085), FALSE | (-0.041, 0.095), FALSE | (-0.313, 0.712), FALSE | (-0.045, 0.108), FALSE | (-0.174, 0.28), FALSE | (-0.02, 0.159), FALSE | (-0.14, 0.116), FALSE |
| CD-cML-S | (-0.052, 0.034), FALSE | (-0.355, 0.172), FALSE | (0.128, 0.176), TRUE | (-0.449, -0.05), TRUE | (0.148, 0.197), TRUE | (-0.071, 0.879), FALSE | (-0.015, 0.073), FALSE | (-0.214, 0.659), FALSE | (-0.019, 0.091), FALSE | (-0.143, 0.207), FALSE | (-0.007, 0.14), FALSE | (-0.126, 0.107), FALSE |
| CD-Ratio-S | (-0.04, 0.037), FALSE | (-0.374, 0.091), FALSE | (0.115, 0.161), TRUE | (-0.198, 0.142), FALSE | (0.139, 0.185), TRUE | (-0.008, 0.346), FALSE | (-0.012, 0.052), FALSE | (-0.233, 0.629), FALSE | (-0.023, 0.084), FALSE | (-0.139, 0.2), FALSE | (-0.011, 0.133), FALSE | (-0.126, 0.107), FALSE |
| CD-Egger-S | (-0.06, 0.059), FALSE | (-0.56, 0.166), FALSE | (0.11, 0.178), TRUE | (-0.432, 0.676), FALSE | (0.134, 0.204), TRUE | (-0.286, 0.676), FALSE | (-0.023, 0.084), FALSE | (-0.257, 0.686), FALSE | (-0.047, 0.12), FALSE | (-0.221, 0.248), FALSE | (-0.049, 0.164), FALSE | (-0.124, 0.108), FALSE |
| MR-cML-DP | (-0.272, 0.235), FALSE | (-0.156, 0.081), FALSE | (0.043, 0.068), TRUE | (-1.33, 0.071), FALSE | (0.03, 0.044), TRUE | (4.405, 11.755), TRUE | (-0.253, 0.634), FALSE | (-0.045, 0.109), FALSE | (-0.093, 0.229), FALSE | (-0.101, 0.142), FALSE | (-0.047, 0.498), FALSE | (-0.036, 0.031), FALSE |
| MR-cML | (-0.198, 0.136), FALSE | (-0.099, 0.046), FALSE | (0.046, 0.064), TRUE | (-1.204, -0.135), TRUE | (0.031, 0.041), TRUE | (6.01, 10.299), TRUE | (-0.069, 0.408), FALSE | (-0.03, 0.101), FALSE | (-0.038, 0.19), FALSE | (-0.081, 0.1), FALSE | (-0.015, 0.46), FALSE | (-0.032, 0.028), FALSE |
| CD-cML-DP | (-0.07, 0.058), FALSE | (-0.568, 0.303), FALSE | (0.118, 0.185), TRUE | (-0.494, 0.026), FALSE | (0.142, 0.209), TRUE | (0.863, 2.463), FALSE | (-0.041, 0.095), FALSE | (-0.313, 0.712), FALSE | (-0.045, 0.108), FALSE | (-0.174, 0.28), FALSE | (-0.02, 0.159), FALSE | (-0.14, 0.116), FALSE |
| CD-cML | (-0.052, 0.034), FALSE | (-0.355, 0.172), FALSE | (0.128, 0.176), TRUE | (-0.448, -0.05), TRUE | (0.148, 0.197), TRUE | (1.259, 2.101), FALSE | (-0.015, 0.073), FALSE | (-0.214, 0.659), FALSE | (-0.019, 0.091), FALSE | (-0.143, 0.207), FALSE | (-0.007, 0.14), FALSE | (-0.126, 0.107), FALSE |
| CD-Ratio | (-0.04, 0.037), FALSE | (-0.374, 0.091), FALSE | (0.115, 0.161), TRUE | (-0.089, 0.243), FALSE | (0.14, 0.186), TRUE | (0.222, 0.548), TRUE | (-0.012, 0.052), FALSE | (-0.233, 0.629), FALSE | (-0.023, 0.084), FALSE | (-0.139, 0.2), FALSE | (-0.011, 0.133), FALSE | (-0.126, 0.107), FALSE |
| CD-Egger | (-0.06, 0.059), FALSE | (-0.56, 0.166), FALSE | (0.11, 0.178), TRUE | (-0.289, 2.059), FALSE | (0.135, 0.206), TRUE | (0.052, 1.643), FALSE | (-0.023, 0.084), FALSE | (-0.257, 0.686), FALSE | (-0.047, 0.12), FALSE | (-0.221, 0.248), FALSE | (-0.049, 0.164), FALSE | (-0.124, 0.108), FALSE |
| Steiger | 0.73, TRUE | 0.143, FALSE | 0.909, TRUE | 0.013, FALSE | 0.907, TRUE | 0.011, FALSE | 0.462, FALSE | 0.038, FALSE | 0.553, TRUE | 0.368, FALSE | 0.633, TRUE | 0.347, FALSE |

Table S5: Inferring causal effects between first 6 risk factors and T2D, in each cell we show the Bonferroni adjusted 1-0.05/48 $\approx$ 0.999 confidence interval of the estimate $\hat{\theta}$ for MR methods, and estimate $\hat{K}$ for CD methods; for Steiger's method, we show proportion of SNPs that give significant result. TRUE/FALSE in each cell indicates whether the result is significant or not, and cells give significant results are marked with red.

| Direction<br>Method | TG to T2D | T2D to TG | LDL to T2D | T2D to LDL | HDL to T2D | T2D to HDL | Height to T2D | T2D to Height | BMI to T2D | T2D to BMI | BF to T2D | T2D to BF |
|---|---|---|---|---|---|---|---|---|---|---|---|---|
| MR-cML-DP-S | (-0.358, 0.459), FALSE | (-0.045, 0.145), FALSE | (-0.348, 0.07), FALSE | (-0.02, 0.043), FALSE | (-0.559, 0.145), FALSE | (-0.08, 0.045), FALSE | (-0.226, 0.134), FALSE | (-0.031, 0.054), FALSE | (0.45, 1.278), TRUE | (-0.103, -0.025), TRUE | (-0.075, 3.161), FALSE | (-0.091, 0.018), FALSE |
| MR-cML-S | (-0.16, 0.27), FALSE | (0.008, 0.107), TRUE | (-0.306, 0.02), FALSE | (-0.014, 0.041), FALSE | (-0.399, -0.009), TRUE | (-0.037, 0.016), FALSE | (-0.176, 0.082), FALSE | (-0.018, 0.041), FALSE | (0.503, 1.158), TRUE | (-0.092, -0.043), TRUE | (0.47, 2.769), TRUE | (-0.071, 0.002), FALSE |
| CD-cML-DP-S | (-0.215, 0.261), FALSE | (-0.054, 0.212), FALSE | (-0.21, 0.038), FALSE | (-0.027, 0.065), FALSE | (-0.315, 0.09), FALSE | (-0.113, 0.06), FALSE | (-0.181, 0.102), FALSE | (-0.037, 0.066), FALSE | (0.402, 1.2), FALSE | (-0.111, -0.023), TRUE | (-0.114, 2.569), FALSE | (-0.117, 0.02), FALSE |
| CD-cML-S | (-0.094, 0.145), FALSE | (-0.003, 0.157), FALSE | (-0.189, 0.008), FALSE | (-0.019, 0.062), FALSE | (-0.217, 0.005), FALSE | (-0.061, 0.025), FALSE | (-0.143, 0.065), FALSE | (-0.021, 0.05), FALSE | (0.453, 1.069), FALSE | (-0.097, -0.045), TRUE | (0.373, 2.216), FALSE | (-0.094, -0.002), TRUE |
| CD-Ratio-S | (-0.096, 0.121), FALSE | (0.012, 0.096), TRUE | (-0.162, 0.027), FALSE | (-0.019, 0.061), FALSE | (-0.193, -0.001), TRUE | (-0.073, 0.007), FALSE | (-0.126, 0.066), FALSE | (-0.023, 0.033), FALSE | (0.427, 1.026), FALSE | (-0.077, -0.031), TRUE | (-0.21, 1.129), FALSE | (-0.083, -0.001), TRUE |
| CD-Egger-S | (-0.155, 0.224), FALSE | (-0.034, 0.178), FALSE | (-0.196, 0.053), FALSE | (-0.017, 0.064), FALSE | (-0.25, 0.038), FALSE | (-0.165, 0.047), FALSE | (-0.171, 0.106), FALSE | (-0.023, 0.045), FALSE | (0.407, 1.16), FALSE | (-0.203, 0.139), FALSE | (-1.205, 2.198), FALSE | (-0.181, 0.107), FALSE |
| MR-cML-DP | (-0.358, 0.459), FALSE | (-0.045, 0.145), FALSE | (-0.348, 0.07), FALSE | (-0.02, 0.043), FALSE | (-0.549, 0.145), FALSE | (-0.08, 0.045), FALSE | (-0.226, 0.134), FALSE | (-0.031, 0.054), FALSE | (0.517, 1.334), TRUE | (-0.103, -0.025), TRUE | (0.48, 3.392), TRUE | (-0.091, 0.018), FALSE |
| MR-cML | (-0.16, 0.27), FALSE | (0.008, 0.107), TRUE | (-0.306, 0.02), FALSE | (-0.014, 0.041), FALSE | (-0.399, -0.009), TRUE | (-0.037, 0.016), FALSE | (-0.176, 0.082), FALSE | (-0.018, 0.041), FALSE | (0.604, 1.204), TRUE | (-0.092, -0.043), TRUE | (1.13, 2.93), TRUE | (-0.071, 0.002), FALSE |
| CD-cML-DP | (-0.215, 0.261), FALSE | (-0.054, 0.212), FALSE | (-0.21, 0.038), FALSE | (-0.027, 0.065), FALSE | (-0.314, 0.097), FALSE | (-0.113, 0.06), FALSE | (-0.181, 0.102), FALSE | (-0.037, 0.066), FALSE | (0.47, 1.252), FALSE | (-0.111, -0.023), TRUE | (0.447, 2.616), FALSE | (-0.117, 0.02), FALSE |
| CD-cML | (-0.094, 0.145), FALSE | (-0.003, 0.157), FALSE | (-0.189, 0.008), FALSE | (-0.019, 0.062), FALSE | (-0.217, 0.005), FALSE | (-0.061, 0.025), FALSE | (-0.143, 0.065), FALSE | (-0.021, 0.05), FALSE | (0.55, 1.115), FALSE | (-0.097, -0.045), TRUE | (0.883, 2.287), FALSE | (-0.094, -0.002), TRUE |
| CD-Ratio | (-0.096, 0.121), FALSE | (0.012, 0.096), TRUE | (-0.162, 0.027), FALSE | (-0.019, 0.061), FALSE | (-0.202, -0.01), TRUE | (-0.073, 0.007), FALSE | (-0.126, 0.066), FALSE | (-0.023, 0.033), FALSE | (0.499, 1.05), FALSE | (-0.077, -0.031), TRUE | (0.282, 1.424), FALSE | (-0.083, -0.001), TRUE |
| CD-Egger | (-0.155, 0.224), FALSE | (-0.034, 0.178), FALSE | (-0.196, 0.053), FALSE | (-0.017, 0.064), FALSE | (-0.282, 0.025), FALSE | (-0.165, 0.047), FALSE | (-0.171, 0.106), FALSE | (-0.023, 0.045), FALSE | (-0.02, 1.356), FALSE | (-0.203, 0.139), FALSE | (-0.383, 2.125), FALSE | (-0.181, 0.107), FALSE |
| Steiger | 0.266, FALSE | 0.172, FALSE | 0.321, FALSE | 0.111, FALSE | 0.348, FALSE | 0.13, FALSE | 0.061, FALSE | 0.034, FALSE | 0, FALSE | 0.171, FALSE | 0, FALSE | 0.6, TRUE |

Table S6: Inferring causal effects between second 6 risk factors and T2D, in each cell we show the Bonferroni adjusted 1-0.05/48 $\approx$ 0.999 confidence interval of the estimate $\hat{\theta}$ for MR methods, and estimate $\hat{K}$ for CD methods; for Steiger's method, we show proportion of SNPs that give significant result. TRUE/FALSE in each cell indicates whether the result is significant or not, and cells give significant results are marked with red.

| Method \ Direction | BW to T2D | T2D to BW | DBP to T2D | T2D to DBP | SBP to T2D | T2D to SBP | FG to T2D | T2D to FG | Smoke to T2D | T2D to Smoke | Alcohol to T2D | T2D to Alcohol |
|---|---|---|---|---|---|---|---|---|---|---|---|---|
| MR-cML-DP-S | (-0.947, 0.193), FALSE | (-0.02, 0.052), FALSE | (-0.007, 0.054), FALSE | (-0.673, 0.382), FALSE | (0.006, 0.038), TRUE | (-0.473, 1.527), FALSE | (0.988, 3.074), TRUE | (0.034, 0.115), TRUE | (-0.535, 0.582), FALSE | (-0.059, 0.027), FALSE | (-0.783, 1.593), FALSE | (-0.043, 0.009), FALSE |
| MR-cML-S | (-0.808, -0.013), TRUE | (-0.013, 0.043), FALSE | (0.002, 0.043), TRUE | (-0.507, 0.026), FALSE | (0.009, 0.033), TRUE | (0.34, 1.047), TRUE | (1.214, 2.835), TRUE | (0.044, 0.104), TRUE | (-0.3, 0.229), FALSE | (-0.056, 0.02), FALSE | (-0.51, 1.319), FALSE | (-0.028, 0), FALSE |
| CD-cML-DP-S | (-0.723, 0.164), FALSE | (-0.026, 0.069), FALSE | (-0.046, 0.433), FALSE | (-0.086, 0.05), FALSE | (0.072, 0.525), TRUE | (-0.027, 0.106), FALSE | (0.437, 1.311), FALSE | (0.084, 0.256), TRUE | (-0.736, 0.797), FALSE | (-0.037, 0.018), FALSE | (-0.609, 1.141), FALSE | (-0.059, 0.015), FALSE |
| CD-cML-S | (-0.605, -0.005), TRUE | (-0.017, 0.057), FALSE | (0.016, 0.347), TRUE | (-0.063, 0.003), FALSE | (0.125, 0.456), TRUE | (0.025, 0.072), TRUE | (0.533, 1.199), FALSE | (0.103, 0.234), TRUE | (-0.457, 0.377), FALSE | (-0.035, 0.014), FALSE | (-0.399, 0.92), FALSE | (-0.036, 0.001), FALSE |
| CD-Ratio-S | (-0.546, 0.002), FALSE | (-0.042, 0.026), FALSE | (0.027, 0.343), TRUE | (-0.009, 0.023), FALSE | (0.14, 0.46), TRUE | (0.025, 0.061), TRUE | (0.228, 0.659), TRUE | (0.11, 0.232), TRUE | (-0.417, 0.36), FALSE | (-0.034, 0.014), FALSE | (-0.42, 0.795), FALSE | (-0.036, -0.002), TRUE |
| CD-Egger-S | (-0.699, 0.176), FALSE | (-0.169, 0.115), FALSE | (-0.022, 0.42), FALSE | (-0.035, 0.054), FALSE | (0.115, 0.527), TRUE | (0.003, 0.102), TRUE | (-0.02, 1.189), FALSE | (0.091, 0.271), TRUE | (-0.488, 0.473), FALSE | (-0.034, 0.015), FALSE | (-0.484, 0.993), FALSE | (-0.047, 0.005), FALSE |
| MR-cML-DP | (-0.996, 0.21), FALSE | (-0.02, 0.052), FALSE | (-0.007, 0.054), FALSE | (-0.673, 0.382), FALSE | (0.005, 0.038), TRUE | (-0.473, 1.527), FALSE | (0.915, 3.254), TRUE | (0.039, 0.108), TRUE | (-0.535, 0.582), FALSE | (-0.059, 0.027), FALSE | (-0.783, 1.593), FALSE | (-0.043, 0.009), FALSE |
| MR-cML | (-0.808, -0.013), TRUE | (-0.013, 0.043), FALSE | (0.002, 0.043), TRUE | (-0.507, 0.026), FALSE | (0.009, 0.033), TRUE | (0.34, 1.047), TRUE | (1.221, 2.827), TRUE | (0.044, 0.104), TRUE | (-0.3, 0.229), FALSE | (-0.056, 0.02), FALSE | (-0.51, 1.319), FALSE | (-0.028, 0), FALSE |
| CD-cML-DP | (-0.771, 0.182), FALSE | (-0.026, 0.069), FALSE | (-0.047, 0.438), FALSE | (-0.086, 0.05), FALSE | (0.074, 0.521), TRUE | (-0.027, 0.106), FALSE | (0.439, 1.399), FALSE | (0.092, 0.246), TRUE | (-0.736, 0.797), FALSE | (-0.037, 0.018), FALSE | (-0.609, 1.141), FALSE | (-0.059, 0.015), FALSE |
| CD-cML | (-0.605, -0.005), TRUE | (-0.017, 0.057), FALSE | (0.016, 0.347), TRUE | (-0.063, 0.003), FALSE | (0.125, 0.456), TRUE | (0.025, 0.072), TRUE | (0.623, 1.305), FALSE | (0.103, 0.234), TRUE | (-0.457, 0.377), FALSE | (-0.035, 0.014), FALSE | (-0.399, 0.92), FALSE | (-0.036, 0.001), FALSE |
| CD-Ratio | (-0.682, -0.145), TRUE | (-0.042, 0.026), FALSE | (0.031, 0.347), TRUE | (-0.009, 0.023), FALSE | (0.158, 0.478), TRUE | (0.025, 0.061), TRUE | (0.28, 0.706), TRUE | (0.119, 0.24), TRUE | (-0.417, 0.36), FALSE | (-0.034, 0.014), FALSE | (-0.42, 0.795), FALSE | (-0.036, -0.002), TRUE |
| CD-Egger | (-1.069, 0.137), FALSE | (-0.169, 0.115), FALSE | (-0.02, 0.449), FALSE | (-0.035, 0.054), FALSE | (0.105, 0.618), TRUE | (0.003, 0.102), TRUE | (-0.444, 1.654), FALSE | (-0.02, 0.429), FALSE | (-0.488, 0.473), FALSE | (-0.034, 0.015), FALSE | (-0.484, 0.993), FALSE | (-0.047, 0.005), FALSE |
| Steiger | 0.038, FALSE | 0.245, FALSE | 0.005, FALSE | 0.067, FALSE | 0.005, FALSE | 0.048, FALSE | 0.091, FALSE | 0.5, TRUE | 0.032, FALSE | 0.355, FALSE | 0, FALSE | 0.359, FALSE |

Table S7: Inferring causal effects between first 6 risk factors and Asthma, in each cell we show the Bonferroni adjusted 1-0.05/48 ≈ 0.999 confidence interval of the estimate $\hat{\theta}$ for MR methods, and estimate $\hat{K}$ for CD methods; for Steiger's method, we show proportion of SNPs that give significant result. TRUE/FALSE in each cell indicates whether the result is significant or not, and cells give significant results are marked with red.

| Direction / Method | TG to Asthma | Asthma to TG | LDL to Asthma | Asthma to LDL | HDL to Asthma | Asthma to HDL | Height to Asthma | Asthma to Height | BMI to Asthma | Asthma to BMI | BF to Asthma | Asthma to BF |
|---|---|---|---|---|---|---|---|---|---|---|---|---|
| MR-cML-DP-S | (-0.291, 0.171), FALSE | (-0.052, 0.073), FALSE | (-0.144, 0.134), FALSE | (-0.061, 0.032), FALSE | (-0.194, 0.154), FALSE | (-0.034, 0.056), FALSE | (-0.082, 0.125), FALSE | (-0.036, 0.022), FALSE | (-0.125, 0.395), FALSE | (-0.032, 0.026), FALSE | (-0.364, 0.617), FALSE | (-0.028, 0.059), FALSE |
| MR-cML-S | (-0.213, 0.085), FALSE | (-0.035, 0.053), FALSE | (-0.123, 0.1), FALSE | (-0.053, 0.029), FALSE | (-0.147, 0.106), FALSE | (-0.03, 0.053), FALSE | (-0.053, 0.117), FALSE | (-0.038, 0.021), FALSE | (-0.08, 0.336), FALSE | (-0.031, 0.024), FALSE | (-0.352, 0.59), FALSE | (-0.025, 0.056), FALSE |
| CD-cML-DP-S | (-0.065, 0.038), FALSE | (-0.161, 0.229), FALSE | (-0.035, 0.034), FALSE | (-0.187, 0.104), FALSE | (-0.046, 0.037), FALSE | (-0.104, 0.174), FALSE | (-0.027, 0.041), FALSE | (-0.11, 0.068), FALSE | (-0.043, 0.148), FALSE | (-0.074, 0.061), FALSE | (-0.105, 0.181), FALSE | (-0.084, 0.17), FALSE |
| CD-cML-S | (-0.048, 0.019), FALSE | (-0.109, 0.166), FALSE | (-0.03, 0.026), FALSE | (-0.158, 0.094), FALSE | (-0.034, 0.025), FALSE | (-0.092, 0.163), FALSE | (-0.017, 0.038), FALSE | (-0.116, 0.066), FALSE | (-0.027, 0.127), FALSE | (-0.073, 0.057), FALSE | (-0.1, 0.174), FALSE | (-0.077, 0.163), FALSE |
| CD-Ratio-S | (-0.052, 0.01), FALSE | (-0.106, 0.139), FALSE | (-0.024, 0.03), FALSE | (-0.179, 0.046), FALSE | (-0.027, 0.029), FALSE | (-0.175, 0.061), FALSE | (-0.025, 0.03), FALSE | (-0.113, 0.066), FALSE | (-0.029, 0.125), FALSE | (-0.073, 0.056), FALSE | (-0.102, 0.171), FALSE | (-0.077, 0.162), FALSE |
| CD-Egger-S | (-0.068, 0.022), FALSE | (-0.166, 0.191), FALSE | (-0.038, 0.04), FALSE | (-0.222, 0.073), FALSE | (-0.038, 0.041), FALSE | (-0.319, 0.168), FALSE | (-0.033, 0.041), FALSE | (-0.108, 0.104), FALSE | (-0.035, 0.132), FALSE | (-0.072, 0.058), FALSE | (-0.122, 0.159), FALSE | (-0.078, 0.162), FALSE |
| MR-cML-DP | (-0.291, 0.171), FALSE | (-0.052, 0.073), FALSE | (-0.144, 0.134), FALSE | (-0.061, 0.032), FALSE | (-0.179, 0.14), FALSE | (-0.034, 0.056), FALSE | (-0.082, 0.125), FALSE | (-0.042, 0.022), FALSE | (-0.125, 0.395), FALSE | (-0.032, 0.026), FALSE | (-0.364, 0.617), FALSE | (-0.028, 0.059), FALSE |
| MR-cML | (-0.213, 0.085), FALSE | (-0.035, 0.053), FALSE | (-0.123, 0.1), FALSE | (-0.053, 0.029), FALSE | (-0.147, 0.106), FALSE | (-0.03, 0.053), FALSE | (-0.053, 0.117), FALSE | (-0.038, 0.021), FALSE | (-0.08, 0.336), FALSE | (-0.031, 0.024), FALSE | (-0.352, 0.59), FALSE | (-0.025, 0.056), FALSE |
| CD-cML-DP | (-0.065, 0.038), FALSE | (-0.161, 0.229), FALSE | (-0.035, 0.034), FALSE | (-0.187, 0.104), FALSE | (-0.043, 0.033), FALSE | (-0.104, 0.174), FALSE | (-0.027, 0.041), FALSE | (-0.127, 0.068), FALSE | (-0.043, 0.148), FALSE | (-0.074, 0.061), FALSE | (-0.105, 0.181), FALSE | (-0.084, 0.17), FALSE |
| CD-cML | (-0.048, 0.019), FALSE | (-0.109, 0.166), FALSE | (-0.03, 0.026), FALSE | (-0.158, 0.094), FALSE | (-0.034, 0.025), FALSE | (-0.092, 0.163), FALSE | (-0.017, 0.038), FALSE | (-0.116, 0.066), FALSE | (-0.027, 0.127), FALSE | (-0.073, 0.057), FALSE | (-0.1, 0.174), FALSE | (-0.077, 0.163), FALSE |
| CD-Ratio | (-0.052, 0.01), FALSE | (-0.106, 0.139), FALSE | (-0.024, 0.03), FALSE | (-0.179, 0.046), FALSE | (-0.029, 0.027), FALSE | (-0.175, 0.061), FALSE | (-0.025, 0.03), FALSE | (-0.127, 0.051), FALSE | (-0.029, 0.125), FALSE | (-0.073, 0.056), FALSE | (-0.102, 0.171), FALSE | (-0.077, 0.162), FALSE |
| CD-Egger | (-0.068, 0.022), FALSE | (-0.166, 0.191), FALSE | (-0.038, 0.04), FALSE | (-0.222, 0.073), FALSE | (-0.065, 0.049), FALSE | (-0.319, 0.168), FALSE | (-0.033, 0.041), FALSE | (-0.437, 0.29), FALSE | (-0.035, 0.132), FALSE | (-0.072, 0.058), FALSE | (-0.122, 0.159), FALSE | (-0.078, 0.162), FALSE |
| Steiger | 0.8, TRUE | 0.2, FALSE | 0.807, TRUE | 0.182, FALSE | 0.845, TRUE | 0.155, FALSE | 0.903, TRUE | 0.024, FALSE | 0.714, TRUE | 0.208, FALSE | 0.346, FALSE | 0.654, TRUE |

Table S8: Inferring causal effects between second 6 risk factors and Asthma, in each cell we show the Bonferroni adjusted 1-0.05/48 $\approx$ 0.999 confidence interval of the estimate $\hat{\theta}$ for MR methods, and estimate $\hat{K}$ for CD methods; for Steiger's method, we show proportion of SNPs that give significant result. TRUE/FALSE in each cell indicates whether the result is significant or not, and cells give significant results are marked with red.

| Method \ Direction | BW to Asthma | Asthma to BW | DBP to Asthma | Asthma to DBP | SBP to Asthma | Asthma to SBP | FG to Asthma | Asthma to FG | Smoke to Asthma | Asthma to Smoke | Alcohol to Asthma | Asthma to Alcohol |
|---|---|---|---|---|---|---|---|---|---|---|---|---|
| MR-cML-DP-S | (-0.213, 0.46), FALSE | (-0.041, 0.022), FALSE | (-0.022, 0.017), FALSE | (-0.425, 0.204), FALSE | (-0.01, 0.011), FALSE | (-0.619, 0.304), FALSE | (-0.692, 0.341), FALSE | (-0.02, 0.048), FALSE | (-0.15, 0.191), FALSE | (-0.05, 0.042), FALSE | (-0.824, 0.569), FALSE | (-0.026, 0.016), FALSE |
| MR-cML-S | (-0.125, 0.379), FALSE | (-0.039, 0.019), FALSE | (-0.017, 0.011), FALSE | (-0.343, 0.057), FALSE | (-0.008, 0.009), FALSE | (-0.517, 0.139), FALSE | (-0.537, 0.164), FALSE | (-0.014, 0.043), FALSE | (-0.157, 0.211), FALSE | (-0.045, 0.038), FALSE | (-0.715, 0.49), FALSE | (-0.021, 0.01), FALSE |
| CD-cML-DP-S | (-0.062, 0.141), FALSE | (-0.135, 0.07), FALSE | (-0.071, 0.054), FALSE | (-0.127, 0.061), FALSE | (-0.054, 0.063), FALSE | (-0.111, 0.054), FALSE | (-0.121, 0.059), FALSE | (-0.117, 0.278), FALSE | (-0.091, 0.117), FALSE | (-0.08, 0.068), FALSE | (-0.245, 0.179), FALSE | (-0.084, 0.052), FALSE |
| CD-cML-S | (-0.036, 0.116), FALSE | (-0.128, 0.061), FALSE | (-0.054, 0.037), FALSE | (-0.102, 0.019), FALSE | (-0.043, 0.049), FALSE | (-0.092, 0.025), FALSE | (-0.094, 0.028), FALSE | (-0.086, 0.251), FALSE | (-0.095, 0.129), FALSE | (-0.073, 0.063), FALSE | (-0.213, 0.157), FALSE | (-0.067, 0.033), FALSE |
| CD-Ratio-S | (-0.044, 0.105), FALSE | (-0.125, 0.062), FALSE | (-0.053, 0.036), FALSE | (-0.076, 0.027), FALSE | (-0.041, 0.049), FALSE | (-0.05, 0.05), FALSE | (-0.081, 0.039), FALSE | (-0.091, 0.23), FALSE | (-0.095, 0.129), FALSE | (-0.072, 0.063), FALSE | (-0.212, 0.157), FALSE | (-0.06, 0.035), FALSE |
| CD-Egger-S | (-0.071, 0.129), FALSE | (-0.126, 0.06), FALSE | (-0.065, 0.049), FALSE | (-0.121, 0.06), FALSE | (-0.05, 0.062), FALSE | (-0.091, 0.104), FALSE | (-0.167, 0.09), FALSE | (-0.131, 0.254), FALSE | (-0.106, 0.125), FALSE | (-0.073, 0.064), FALSE | (-0.247, 0.185), FALSE | (-0.072, 0.057), FALSE |
| MR-cML-DP | (-0.213, 0.46), FALSE | (-0.041, 0.022), FALSE | (-0.022, 0.017), FALSE | (-0.425, 0.204), FALSE | (-0.01, 0.011), FALSE | (-0.619, 0.304), FALSE | (-0.692, 0.341), FALSE | (-0.02, 0.048), FALSE | (-0.15, 0.191), FALSE | (-0.05, 0.042), FALSE | (-0.824, 0.569), FALSE | (-0.026, 0.016), FALSE |
| MR-cML | (-0.125, 0.379), FALSE | (-0.039, 0.019), FALSE | (-0.017, 0.011), FALSE | (-0.343, 0.057), FALSE | (-0.008, 0.009), FALSE | (-0.517, 0.139), FALSE | (-0.537, 0.164), FALSE | (-0.014, 0.043), FALSE | (-0.157, 0.211), FALSE | (-0.045, 0.038), FALSE | (-0.715, 0.49), FALSE | (-0.021, 0.01), FALSE |
| CD-cML-DP | (-0.062, 0.141), FALSE | (-0.135, 0.07), FALSE | (-0.071, 0.054), FALSE | (-0.127, 0.061), FALSE | (-0.054, 0.063), FALSE | (-0.111, 0.054), FALSE | (-0.121, 0.059), FALSE | (-0.117, 0.278), FALSE | (-0.091, 0.117), FALSE | (-0.08, 0.068), FALSE | (-0.245, 0.179), FALSE | (-0.084, 0.052), FALSE |
| CD-cML | (-0.036, 0.116), FALSE | (-0.128, 0.061), FALSE | (-0.054, 0.037), FALSE | (-0.102, 0.019), FALSE | (-0.043, 0.049), FALSE | (-0.092, 0.025), FALSE | (-0.094, 0.028), FALSE | (-0.086, 0.251), FALSE | (-0.095, 0.129), FALSE | (-0.073, 0.063), FALSE | (-0.213, 0.157), FALSE | (-0.067, 0.033), FALSE |
| CD-Ratio | (-0.044, 0.105), FALSE | (-0.125, 0.062), FALSE | (-0.053, 0.036), FALSE | (-0.076, 0.027), FALSE | (-0.041, 0.049), FALSE | (-0.05, 0.05), FALSE | (-0.081, 0.039), FALSE | (-0.091, 0.23), FALSE | (-0.095, 0.129), FALSE | (-0.072, 0.063), FALSE | (-0.212, 0.157), FALSE | (-0.06, 0.035), FALSE |
| CD-Egger | (-0.071, 0.129), FALSE | (-0.126, 0.06), FALSE | (-0.065, 0.049), FALSE | (-0.121, 0.06), FALSE | (-0.05, 0.062), FALSE | (-0.091, 0.104), FALSE | (-0.167, 0.09), FALSE | (-0.131, 0.254), FALSE | (-0.106, 0.125), FALSE | (-0.073, 0.064), FALSE | (-0.247, 0.185), FALSE | (-0.072, 0.057), FALSE |
| Steiger | 0.712, TRUE | 0.271, FALSE | 0.5, TRUE | 0.024, FALSE | 0.513, TRUE | 0.031, FALSE | 0.414, FALSE | 0.517, TRUE | 0.471, FALSE | 0.5, TRUE | 0.357, FALSE | 0.405, TRUE |

#### S2.2 Pairs of 4 Diseases

Figure S16: Causal relationship between pairs of 4 diseases.

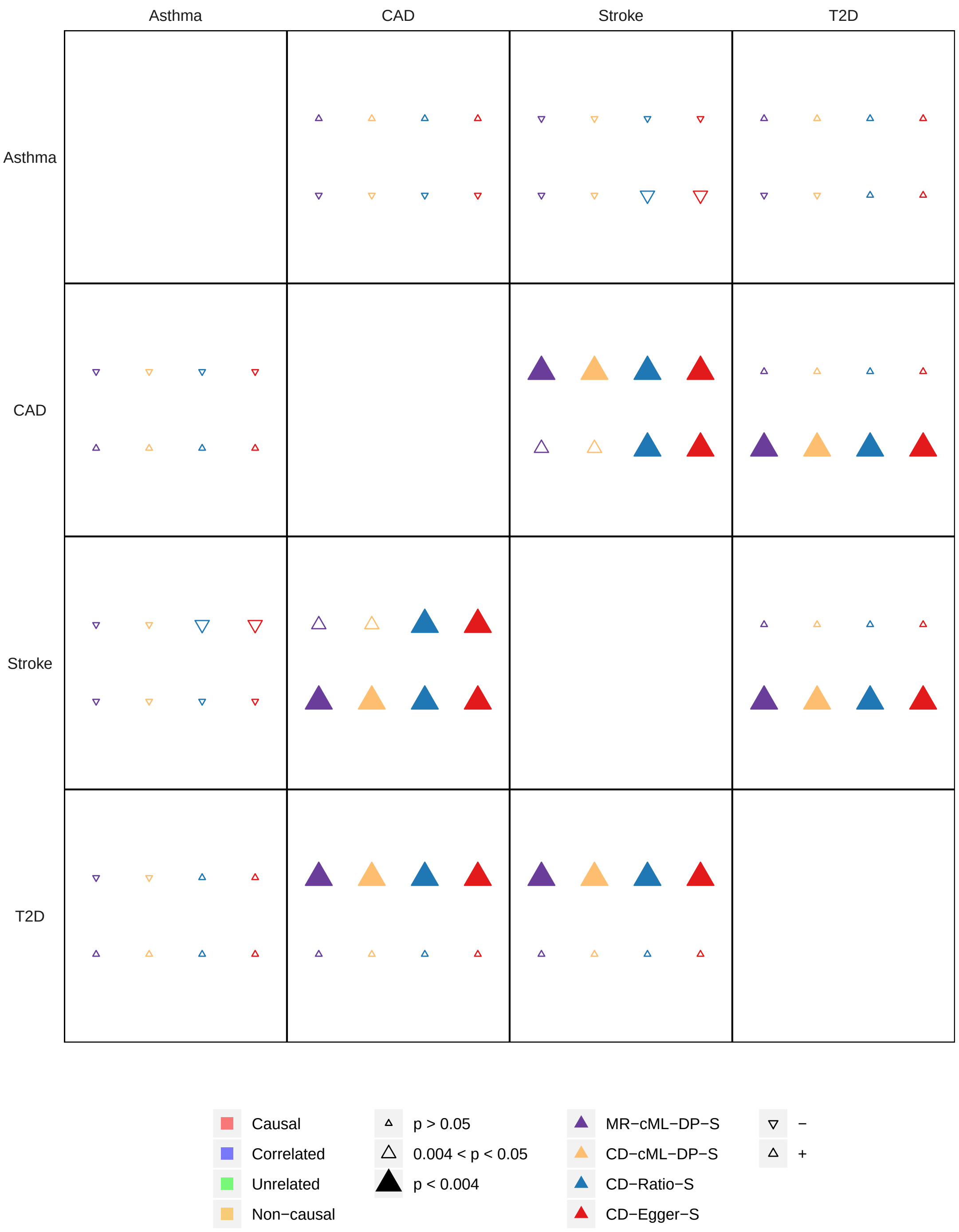

Table S9: Inferring causal effects between pairs of 4 diseases, in each cell we show the Bonferroni adjusted 1-0.05/48 $\approx$ 0.999 confidence interval of the estimate $\hat{\theta}$ for MR methods, and estimate $\hat{K}$ for CD methods; for Steiger's method, we show proportion of SNPs that give significant result. TRUE/FALSE in each cell indicates whether the result is significant or not, and cells give significant results are marked with red.

| Method \ Direction | CAD to Stroke | Stroke to CAD | CAD to T2D | T2D to CAD | CAD to Asthma | Asthma to CAD | Stroke to T2D | T2D to Stroke | Stroke to Asthma | Asthma to Stroke | T2D to Asthma | Asthma to T2D |
|---|---|---|---|---|---|---|---|---|---|---|---|---|
| MR-cML-DP-S | (0.102, 0.318), TRUE | (-0.032, 0.414), FALSE | (-0.221, 0.233), FALSE | (0.009, 0.134), TRUE | (-0.179, 0.17), FALSE | (-0.043, 0.099), FALSE | (-0.313, 0.523), FALSE | (0.011, 0.144), TRUE | (-0.442, 0.132), FALSE | (-0.077, 0.044), FALSE | (-0.126, 0.095), FALSE | (-0.209, 0.233), FALSE |
| MR-cML-S | (0.135, 0.25), TRUE | (0.05, 0.309), TRUE | (-0.094, 0.165), FALSE | (0.019, 0.111), TRUE | (-0.125, 0.13), FALSE | (-0.019, 0.084), FALSE | (-0.223, 0.473), FALSE | (0.023, 0.124), TRUE | (-0.397, 0.068), FALSE | (-0.069, 0.04), FALSE | (-0.096, 0.072), FALSE | (-0.117, 0.145), FALSE |
| CD-cML-DP-S | (0.099, 0.29), TRUE | (-0.022, 0.432), FALSE | (-0.592, 0.623), FALSE | (0.003, 0.048), TRUE | (-0.196, 0.175), FALSE | (-0.039, 0.087), FALSE | (-0.878, 1.387), FALSE | (0.003, 0.048), TRUE | (-0.511, 0.174), FALSE | (-0.065, 0.035), FALSE | (-0.049, 0.04), FALSE | (-0.496, 0.565), FALSE |
| CD-cML-S | (0.131, 0.232), TRUE | (0.057, 0.323), TRUE | (-0.278, 0.449), FALSE | (0.007, 0.039), TRUE | (-0.139, 0.095), FALSE | (-0.017, 0.074), FALSE | (-0.674, 1.276), FALSE | (0.007, 0.041), TRUE | (-0.449, 0.097), FALSE | (-0.058, 0.032), FALSE | (-0.037, 0.03), FALSE | (-0.281, 0.347), FALSE |
| CD-Ratio-S | (0.124, 0.219), TRUE | (0.088, 0.322), TRUE | (-0.278, 0.385), FALSE | (0.009, 0.037), TRUE | (-0.141, 0.06), FALSE | (-0.025, 0.058), FALSE | (-0.44, 1.175), FALSE | (0.006, 0.039), TRUE | (-0.401, 0.043), FALSE | (-0.058, 0.032), FALSE | (-0.03, 0.03), FALSE | (-0.277, 0.305), FALSE |
| CD-Egger-S | (0.108, 0.271), TRUE | (0.002, 0.525), TRUE | (-0.47, 0.636), FALSE | (0.005, 0.044), TRUE | (-0.176, 0.111), FALSE | (-0.071, 0.142), FALSE | (-0.784, 1.365), FALSE | (0.006, 0.042), TRUE | (-0.535, 0.078), FALSE | (-0.058, 0.034), FALSE | (-0.043, 0.046), FALSE | (-0.355, 0.382), FALSE |
| MR-cML-DP | (0.098, 0.327), TRUE | (-0.021, 0.417), FALSE | (-0.221, 0.233), FALSE | (0.009, 0.134), TRUE | (-0.178, 0.165), FALSE | (-0.043, 0.099), FALSE | (-0.313, 0.523), FALSE | (0.011, 0.144), TRUE | (-0.442, 0.132), FALSE | (-0.077, 0.044), FALSE | (-0.126, 0.095), FALSE | (-0.209, 0.233), FALSE |
| MR-cML | (0.135, 0.25), TRUE | (0.05, 0.309), TRUE | (-0.094, 0.165), FALSE | (0.019, 0.111), TRUE | (-0.125, 0.13), FALSE | (-0.019, 0.084), FALSE | (-0.223, 0.473), FALSE | (0.023, 0.124), TRUE | (-0.397, 0.068), FALSE | (-0.069, 0.04), FALSE | (-0.096, 0.072), FALSE | (-0.117, 0.145), FALSE |
| CD-cML-DP | (0.093, 0.303), TRUE | (-0.028, 0.456), FALSE | (-0.592, 0.623), FALSE | (0.003, 0.048), TRUE | (-0.199, 0.173), FALSE | (-0.039, 0.087), FALSE | (-0.878, 1.387), FALSE | (0.003, 0.048), TRUE | (-0.511, 0.174), FALSE | (-0.065, 0.035), FALSE | (-0.049, 0.04), FALSE | (-0.496, 0.565), FALSE |
| CD-cML | (0.131, 0.232), TRUE | (0.057, 0.323), TRUE | (-0.278, 0.449), FALSE | (0.007, 0.039), TRUE | (-0.139, 0.095), FALSE | (-0.017, 0.074), FALSE | (-0.674, 1.276), FALSE | (0.007, 0.041), TRUE | (-0.449, 0.097), FALSE | (-0.058, 0.032), FALSE | (-0.037, 0.03), FALSE | (-0.281, 0.347), FALSE |
| CD-Ratio | (0.131, 0.225), TRUE | (0.123, 0.353), TRUE | (-0.278, 0.385), FALSE | (0.009, 0.037), TRUE | (-0.139, 0.061), FALSE | (-0.025, 0.058), FALSE | (-0.44, 1.175), FALSE | (0.006, 0.039), TRUE | (-0.401, 0.043), FALSE | (-0.058, 0.032), FALSE | (-0.03, 0.03), FALSE | (-0.277, 0.305), FALSE |
| CD-Egger | (0.115, 0.293), TRUE | (0.001, 0.61), TRUE | (-0.47, 0.636), FALSE | (0.005, 0.044), TRUE | (-0.257, 0.128), FALSE | (-0.071, 0.142), FALSE | (-0.784, 1.365), FALSE | (0.006, 0.042), TRUE | (-0.535, 0.078), FALSE | (-0.058, 0.034), FALSE | (-0.043, 0.046), FALSE | (-0.355, 0.382), FALSE |
| Steiger | 0.768, TRUE | 0.146, FALSE | 0.014, FALSE | 0.25, FALSE | 0.411, TRUE | 0.233, FALSE | 0, FALSE | 0.5, FALSE | 0.312, FALSE | 0.531, TRUE | 0.414, FALSE | 0, FALSE |
